# Molecular Connectomics Reveals a Glucagon-Like Peptide 1 Sensitive Neural Circuit for Satiety

**DOI:** 10.1101/2023.10.31.564990

**Authors:** Addison N. Webster, Jordan J. Becker, Chia Li, Dana C. Schwalbe, Damien Kerspern, Eva O. Karolczak, Catherine Bundon, Roberta A. Onoharigho, Maisie Crook, Maira Jalil, Elizabeth N. Godschall, Emily G. Dame, Adam Dawer, Dylan Matthew Belmont-Rausch, Tune H. Pers, Andrew Lutas, Naomi Habib, Ali D. Güler, Michael J. Krashes, John N. Campbell

## Abstract

Liraglutide and other agonists of the glucagon-like peptide 1 receptor (GLP-1RAs) are effective weight loss drugs, but how they suppress appetite remains unclear. One potential mechanism is by activating neurons which inhibit hunger-promoting Agouti-related peptide (AgRP) neurons of the arcuate hypothalamus (Arc). To identify these afferents, we developed a method combining rabies-based connectomics with single-nuclei transcriptomics. Applying this method to AgRP neurons predicted at least 21 afferent subtypes in the mouse mediobasal and paraventricular hypothalamus. Among these are *Trh+* Arc neurons (Trh^Arc^), inhibitory neurons which express the *Glp1r* gene and are activated by the GLP-1RA liraglutide. Activating Trh^Arc^ neurons inhibits AgRP neurons and feeding in an AgRP neuron-dependent manner. Silencing Trh^Arc^ neurons causes over-eating and weight gain and attenuates liraglutide’s effect on body weight. Our results demonstrate a widely applicable method for molecular connectomics, comprehensively identify local inputs to AgRP neurons, and reveal a circuit through which GLP-1RAs suppress appetite.

## INTRODUCTION

Liraglutide, a clinically approved agonist of the glucagon-like peptide 1 receptor (GLP-1RA), potently suppresses appetite through its action on unidentified neurons ^1–5^. GLP-1Rs are abundantly expressed by vagal sensory neurons in the periphery, the dorsal vagal complex of the medulla, and the arcuate nucleus of the hypothalamus (Arc) ^6,7^. However, GLP-1Rs on vagal sensory neurons and *Phox2b*-expressing neurons of the dorsal vagal complex are not required for liraglutide to suppress appetite ^2,3^ (but see also reference ^8^). On the other hand, antagonizing GLP-1Rs in the Arc blocks the appetite-suppressing effects of liraglutide ^3^. Fluorescently labeled liraglutide binds to Arc neurons after systemic administration ^3^, likely due to its transport by tanycytes ^9^, specialized ependymal cells along the third ventricle. Thus, liraglutide can directly act on Arc neurons to suppress appetite.

Agouti-related peptide (AgRP) neurons are a conserved, molecularly distinct neuronal population that play a critical role in energy balance ^4,10,11^. Activating AgRP neurons rapidly and robustly drives feeding behavior ^12–14^. While acute liraglutide injection fails to alter *in vivo* AgRP neuronal activity over the course of minutes ^15^, 24 hour pre-treatment strongly blunts the response of AgRP neurons to food ^16^. Furthermore, liraglutide can indirectly inhibit hunger-promoting AgRP neurons ^3,10,16,17^, providing a potential anorectic mechanism behind this medication. Electrophysiological recordings demonstrated that liraglutide hyperpolarizes and reduces the firing of AgRP cells by increasing inhibitory tone ^3,16^. Other studies have identified local GABAergic input to AgRP neurons ^18^, though the neurons providing this input, and whether they express GLP-1R, remain unknown. Here, by combining rabies-based connectomics with single-cell transcriptomics, we identified a liraglutide-activated subtype of Arc neuron that inhibits AgRP neurons to suppress hunger. Moreover, we found that the activity of these neurons contributes to both the appetite-suppressing effects of liraglutide and the regulation of body weight and food intake.

## RESULTS

### RAMPANT Identifies Transcriptionally Distinct Afferents to AgRP Neurons

To identify which neurons synapse on AgRP neurons, we developed a method for molecular connectomics, RAMPANT (Rabies Afferent Mapping by Poly-A Nuclear Transcriptomics), and applied it to AgRP neurons. We targeted rabies infection to AgRP neurons and their presynaptic partners by injecting a “helper” adeno-associated virus (AAV) that Cre-dependently expresses the avian TVA (tumor virus A) receptor and an optimized rabies glycoprotein (oG) into the Arc of Agrp-Cre mice (n = 41 mice, Fig. 1A). The TVA receptor permits entry of Envelope A (EnvA)-pseudotyped viruses such as [EnvA]-rabies ^19^ into cells, while the rabies glycoprotein is necessary for the virus then to spread trans-synaptically ^20,21^. We later injected the Arc of the same mice with an EnvA-pseudotyped, glycoprotein-deficient rabies virus expressing a nuclear-localized fluorescent protein, H2b-mCherry, [EnvA]rabies-H2b-mCherry (Fig. 1A). Importantly, injecting the same series of helper AAV and rabies viruses into the brains of wildtype C57BL/6j mice failed to label any cells with either GFP or H2b-mCherry, validating the specificity of these viruses (Fig. S1A; n=5 mice).

**Figure 1.**
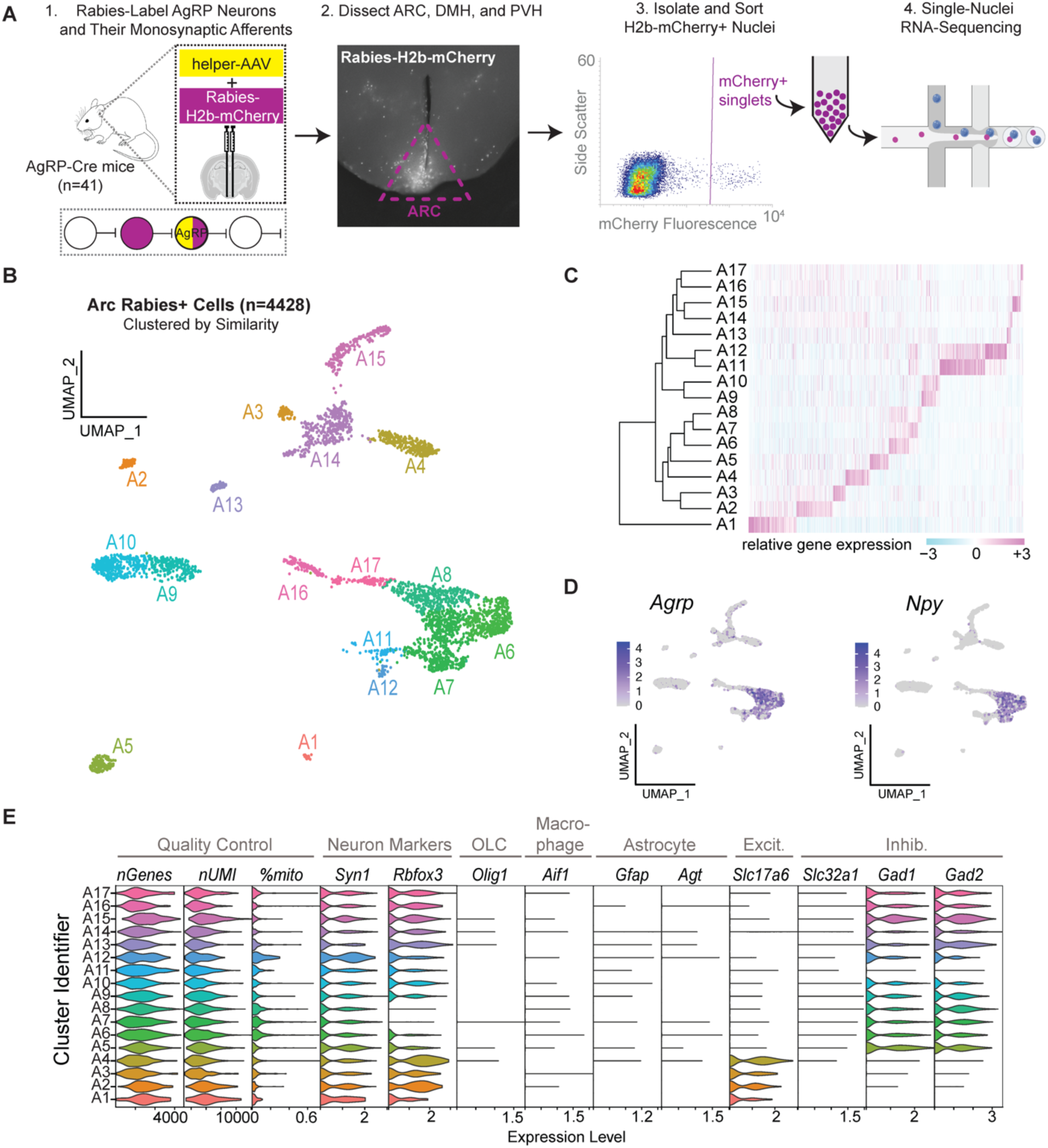
– RAMPANT Identifies Transcriptionally Distinct Afferents To AgRP Neurons. **A**, Schematic of RAMPANT method. **B,** UMAP (uniform manifold approximation and projection) plot of 4,428 rabies+ cell nuclei from the arcuate hypothalamus, clustered by expression of high-variance genes and colored according to cluster identity. **C,** (left) Dendrogram illustrating relatedness of 17 cell clusters; (right) heatmap visualizing expression of genes significantly enriched in each cell cluster. **D,** UMAP recolored to illustrate expression levels of the AgRP neuron marker genes, *Agrp* and *Npy*. **E,** Violin plots showing expression of the following: quality control metrics (number of genes, nGenes; number of unique molecular identifiers, nUMI; percentage of reads from mitochondrial genes, %mito); neuronal marker genes (*Syn1*, NeuN/*Rbfox3*); oligodendrocyte marker gene (*Olig1*); macrophage marker gene (*Aif1*); astrocyte marker genes (*Gfap*, *Agt*); excitatory neuron marker gene (*Slc17a6*); and inhibitory neuron marker genes (*Slc32a1*, *Gad1*, *Gad2*).

To molecularly identify rabies-infected afferents to AgRP neurons in the Arc, dorsomedial hypothalamus (DMH) and paraventricular hypothalamus (PVH), we isolated H2b-mCherry+ cell nuclei from each of these regions and profiled their genome-wide mRNA content by single-nuclei RNA-sequencing (snRNA-seq; Fig. 1A). We focused on these brain regions since they contain the highest densities of cells labeled by rabies via AgRP neurons (Fig. S1B-C) ^22,23^ and because the Arc and DMH are potential sources of Glp1r+ synaptic input to AgRP neurons (Fig. S1D). We chose to profile nuclei instead of whole cells because enzymatic dissociation appears to deplete rabies-infected neurons, potentially due to poor cell viability ^24^, and because a cell’s nuclear RNA profile is sufficient to identify its molecular type ^25–27^. For control samples, we similarly profiled AgRP neurons labeled instead with a Cre-dependent AAV expressing H2b-mCherry (AAV-DIO-H2b-mCherry). Our initial dataset (“all rabies”) of rabies-labeled cells from the Arc, DMH, and PVH totaled 5,770 cells and averaged 2,296 +/− 642 genes detected per cell, after filtering for quality control (mean +/− st. dev.; (Fig. S2, S3A). Grouping these cells based on their expression of high-variance genes yielded 24 cell clusters (Fig. S3B,C), which we annotated by brain region of origin (Fig. S3D). Some of the clusters contained cells from multiple brain regions, suggesting that these cells may reside at or across the regional borders approximated during tissue dissection (Fig. S3E). Importantly, among the clusters were two neuron subtypes, which likely correspond to known presynaptic partners of AgRP neurons: PVH cells expressing the gene *Trh*, which encodes thyrotropin-releasing hormone (Fig. S3F) ^23^, and DMH cells expressing the leptin receptor gene, *Lepr*, and *Glp1r* (Fig. S3G,H) ^18,28^.

We focused our analysis on the Arc, since it contains many afferents to AgRP neurons ^22,23^, potentially including glucagon-like peptide (GLP1)– and leptin-sensing populations which could inhibit AgRP neurons to suppress hunger ^3,18^. To identify Arc cell subtypes, we subsetted Arc cells and re-clustered them apart from non-Arc cells. Filtering out likely cell doublets and low-quality transcriptomes left 4,428 cells, in which we detected 2,308 +/− 634 genes per cell (mean +/− st. dev.). Grouping these cells based on high-variance transcripts yielded 17 molecularly distinct clusters (Fig. 1B-C). As expected, some of these clusters showed enriched expression of AgRP neuron marker genes (*i.e.*, *Agrp*, *Npy*; Fig. 1D). We identified these clusters as neurons based on their content of neuron marker transcripts *Syn1* and *Rbfox3* and relative lack of glial and stromal cell marker transcripts (Fig 1E). Based on their expression of the neurotransmitter phenotype genes *Slc17a6*, *Gad1*, and *Gad2*, we classified 4 of the clusters as glutamatergic, 12 as GABAergic, and 1 as molecularly ambiguous (Fig. 1E). Of note, the inhibitory neuron marker gene, *Slc32a1*, which encodes the vesicular GABA transporter (vGAT), was barely detectable among the *Gad1*+/*Gad2*+ clusters (Fig. 1E). This is consistent with previous reports that rabies infection downregulates *Slc32a1* expression ^29^. Together, our findings demonstrate that RAMPANT can identify transcriptionally distinct groups of neurons.

### Most Marker Genes of AgRP Neurons Are Not Significantly Affected by Rabies Infection

Rabies infection can alter expression of genes that distinguish molecular cell types, such as *Slc32a1*, which represents a potential confound to identifying cell types ^24,29^. For instance, if rabies downregulates genes normally enriched in a cell type (positive markers) or upregulates genes normally found in neighboring cell types (negative markers), this could undermine the ability to match a rabies-infected cell with molecular cell types from a reference atlas.

To investigate the potential confound of rabies-altered gene expression, we compared the transcriptomes of individual AgRP neurons infected with either with the helper AAV and [EnvA]rabies-H2b-mCherry (AAV+rabies), or just AAV-DIO-H2b-mCherry alone (AAV-only; n=689 AAV+rabies cells and 2,133 AAV-only cells). We assigned a neuronal subtype identity to each AAV-only or AAV+rabies cell by transferring cell-type labels from a reference dataset of molecular neuron subtypes in the Arc-ME ^30^ (Fig. 2A). This label-transfer method predicted both populations of cells, AAV+rabies and AAV-only, to be AgRP neurons with similarly high confidence (Fig. 2B).

**Figure 2.**
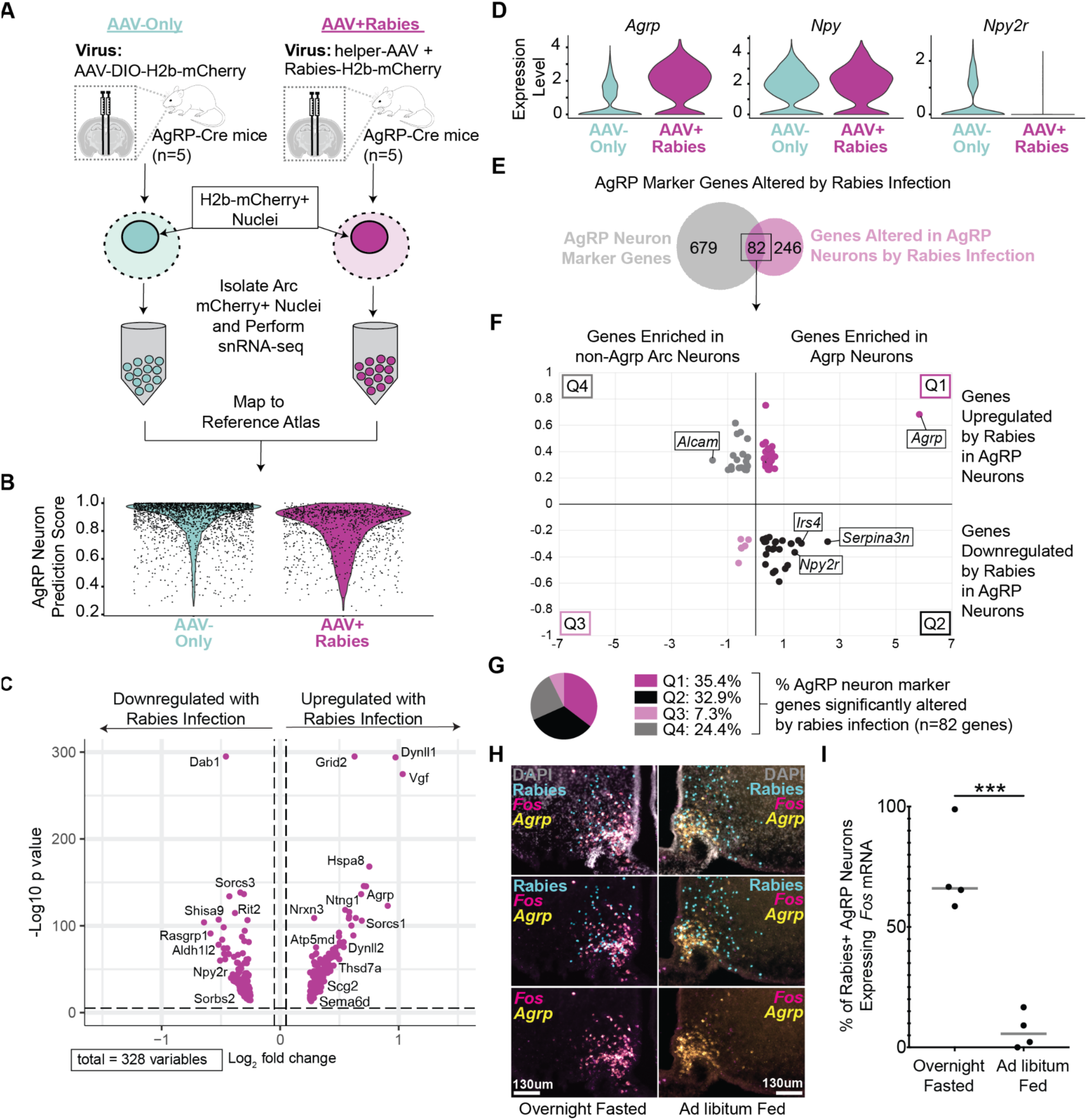
– Molecular and Functional Features of AgRP Neurons Five Days After Rabies Infection. **A**, Schematic for comparing gene expression between AgRP neurons infected with AAV-only or with AAV and rabies (AAV+rabies). **B,** Violin plots comparing prediction scores for AAV-only cells *vs*. AAV+rabies cells mapping to the AgRP neuron cluster from the reference dataset ^30^. **C,** Volcano plot of genes differentially expressed to a significant extent between AAV-only AgRP neurons and AAV+Rabies AgRP neurons. **D,** Violin plots comparing AAV-only AgRP neurons and AAV+Rabies AgRP neurons in terms of their expression of the AgRP neuron marker genes *Agrp, Npy,* and *Npy2r*. **E,** Venn diagram showing overlap of AgRP neuron marker genes (positive and negative) and genes significantly altered by rabies in AgRP neurons. **F,** For each AgRP neuron marker gene altered by rabies, its enrichment in AgRP neurons and non-AgRP Arc neurons compared to its alteration by rabies infection in AgRP neurons. For instance, the top right quadrant (Q1) contains genes significantly enriched in AgRP neurons which are upregulated by rabies. **G,** Pie graph of the number of genes per quadrant in Figure 2F as a percentage of the total number of AgRP neuron marker genes significantly affected by rabies. **H,** Representative images of *Agrp* and *Fos* RNA FISH in rabies-infected Arc cells after overnight fasting or ad libitum feeding. **I,** Comparison of *Fos* RNA+ rabies-infected AgRP neurons between fed and fasted mice (n=4); unpaired Student’s t test, t=6.701, df=6; *** p<0.001.

Comparing the AAV+rabies AgRP neurons to the AAV-only AgRP neurons revealed 328 genes as significantly altered by rabies infection (Fig. 2C). Among these were genes which distinguish AgRP neurons from other Arc neurons in our reference atlas ^30^. For instance, among positive markers for AgRP neurons, rabies infection upregulated *Agrp* and downregulated *Npy2r*, though expression of *Npy* did not change significantly (Fig. 2D). Importantly, only 82 of the 328 rabies-affected genes (25%) were AgRP neuron marker genes, and these represented 11% of the AgRP neuron marker genes (n=761 positive and negative markers combined; Fig. 2E). Of these 82 rabies-affected marker genes for AgRP neurons, 35 (42.7%) were altered in the same direction as their enrichment in AgRP neurons – e.g., 29 genes enriched in AgRP neurons, positive markers, were upregulated by rabies in AgRP neurons (Fig. 2F,G). Our results therefore demonstrate that most of the positive and negative marker genes for AgRP neurons are not significantly affected by rabies five days after infection. Among the AgRP neuron marker genes altered by rabies, roughly half were altered in the same direction as their enrichment in AgRP neurons and so may be less likely to confound cell-type identification. These results suggest that cells acutely infected with rabies can still be identified based on their expression of most cell-type marker genes.

To determine whether rabies infection alters AgRP neuron functionality, we assessed their expression of the activation marker and immediate early gene, *Fos*, after fasting. Specifically, we analyzed *Agrp* and *Fos* co-expression in rabies-infected Arc neurons by RNA fluorescence in situ hybridization (RNA FISH) after an overnight fast or ad libitum feeding. Our results show that fasting significantly increases the percentage of *Fos* RNA+ AgRP neurons, relative to AgRP neurons in ad libitum fed mice (Fig. 2H-I). Thus, fasting activates rabies-infected AgRP neurons as it does AgRP neurons not infected with rabies ^31,32^.

### Label Transfer Predicts Molecular Subtypes of Rabies-Infected Arc Neurons

We assigned a neuronal subtype identity to each Arc rabies+ cells by transferring cell-type labels from a reference dataset of molecular neuron subtypes in the Arc-ME ^28^ (Fig. 3A,B). Removing rabies-infected cells that were identified with low confidence (cell-type prediction score <0.5 out of 1) left 3,593 cells (81%) for further analysis. Our results show that the Arc rabies-infected cells represent 19 of the 31 Arc-ME neuron subtypes in our reference dataset, including AgRP neurons as expected (Fig. 3C,D). Of these 19 subtypes, 14 originated from the Arc, whereas four likely came from neighboring regions, such as the ventromedial hypothalamus (VMH) and retrochiasmatic area ^30^ (Fig. 3C,D), as an artifact of dissection. Several of the 14 Arc subtypes were over-represented in our Arc RAMPANT dataset relative to the reference, including n11.Trh/Cxcl12 neurons, n21.Pomc/Glipr1 neurons, n20.Kiss1/Tac2 neurons, and n08.Th/Slc6a3 neurons (Fig. 3D), consistent with non-random sampling of Arc cells. In addition, we identified Arc neuron subtypes which likely correspond to previously identified afferents to AgRP neurons: n20.Kiss1/Tac2 neurons (Kisspeptin, Neurokinin B/Tac2, Dynorphin, or KNDy, neurons) ^33^; n08.Th/Slc6a3 neurons (tuberoinfundibular, or TIDA, neurons) ^34^; and multiple subtypes of *Drd1*+, *Lepr*+, and *Glp1r*+ neurons ^3,18,35^ (Fig. S4A). Thus, our RAMPANT dataset includes all known afferents to AgRP neurons and predicts many additional subtypes of afferent neurons within the Arc.

**Figure 3.**
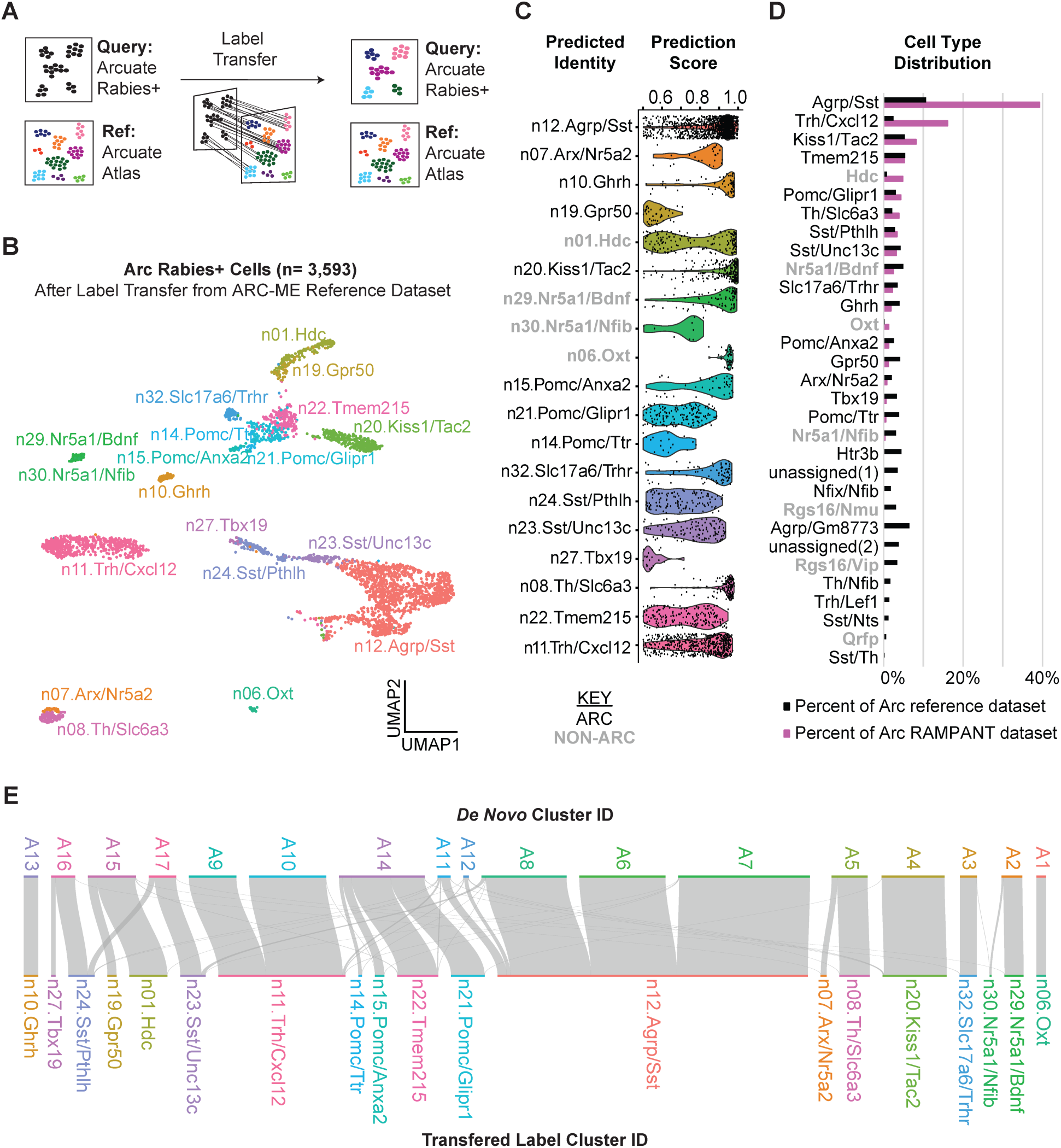
– Molecular Subtypes of Rabies-Infected Arc Neurons Predicted by Label Transfer. **A**, Schematic of method for transferring labels from a reference single-cell RNA-seq dataset to rabies+ cells. **B,** UMAP plot of 3,593 Arc rabies+ cell transcriptomes clustered by expression of highly variable genes and colored by cell-type label transferred from a reference transcriptomic atlas of Arc neuron subtypes ^30^. **C,** Violin plots of cell type prediction scores from mapping rabies+ cells to the reference atlas dataset ^30^. **D,** Distribution of rabies+ Arc neurons and reference Arc neurons across Arc neuron subtypes. **E,** River plot of relationship between *de novo* cell clusters and reference-labeled cell clusters.

In some cases, we observed that neurons from the same de novo cluster mapped to different reference clusters. For example, de novo cluster A14 consisted of neurons which mapped to the n21.Pomc/Glipr1, n22.Tmem215, n15.Pomc/Anxa2, and n14.Pomc/Ttr subtypes (Fig. 3E). This observation confirms previous reports that reference-guided cell annotation can help separate closely related cell types more efficiently than de novo clustering ^36^. Together, our results demonstrate that rabies-infected neurons can be identified by comparing them to uninfected neurons, despite the transcriptional artifacts of rabies infection.

To validate our results against a more comprehensive reference atlas, we repeated the label transfer method but used a consensus cell-type atlas of the mouse hypothalamus, the mouse HypoMap ^37^. By transferring HypoMap cell-type labels to our Arc rabies+ cells, we detected 25 different neuronal subtypes (Fig. S4B-C), more than the 19 neuronal subtypes detected with our Arc-ME reference atlas (correspondence between label transfer results shown in Fig. S4D). We also extended this approach to our all-rabies dataset and found 56 cell types and 14 hypothalamic brain regions represented (Fig. S5A-D). Thus, using a comprehensive, consensus-based reference of molecular cell types can improve RAMPANT’s precision in predicting cell types and their brain region of origin.

### RNA Fluorescence *In Situ* Hybridization Validates RAMPANT Neuron Subtype Predictions

Our RAMPANT study predicts at least 14 molecular subtypes of Arc neurons labeled by rabies via AgRP neurons. To validate these predictions, we used RNA FISH staining in tissue following monosynaptic rabies-H2b-mCherry tracing from AgRP neurons. We selected four Arc neuron subtypes, each marked by its expression of *Slc6a3*, *Ghrh*, *Pomc,* or *Trh* (Fig. 4A). We validated these subtypes by co-localizing rabies-H2b-mCherry with fluorescently labeled transcripts of each subtype-specific marker by RNA FISH (Fig. 4B). Quantifying the rate of colocalization for each subtype indicates that *Slc6a3+* neurons, *Ghrh+* neurons, *Pomc+* neurons, and *Trh+* neurons constitute 4.56%, 2.98%, 6.61%, and 16.16% of rabies-labeled afferents to AgRP neurons in the Arc, respectively (n=3 mice per subtype; Fig. 4C). These estimates correlate highly with the percentages of corresponding cell types in our RAMPANT dataset: n08.Th/Slc6a3 neurons, n10.Ghrh neurons, Pomc neurons (all subtypes combined), and n11.Trh/Cxcl12 neurons comprise 3.84%, 1.84%, 5.93%, and 16.06% of the cells (Fig. 4D). Thus, our results validate RAMPANT’s predictions of these subtype identities and their proportions.

**Figure 4.**
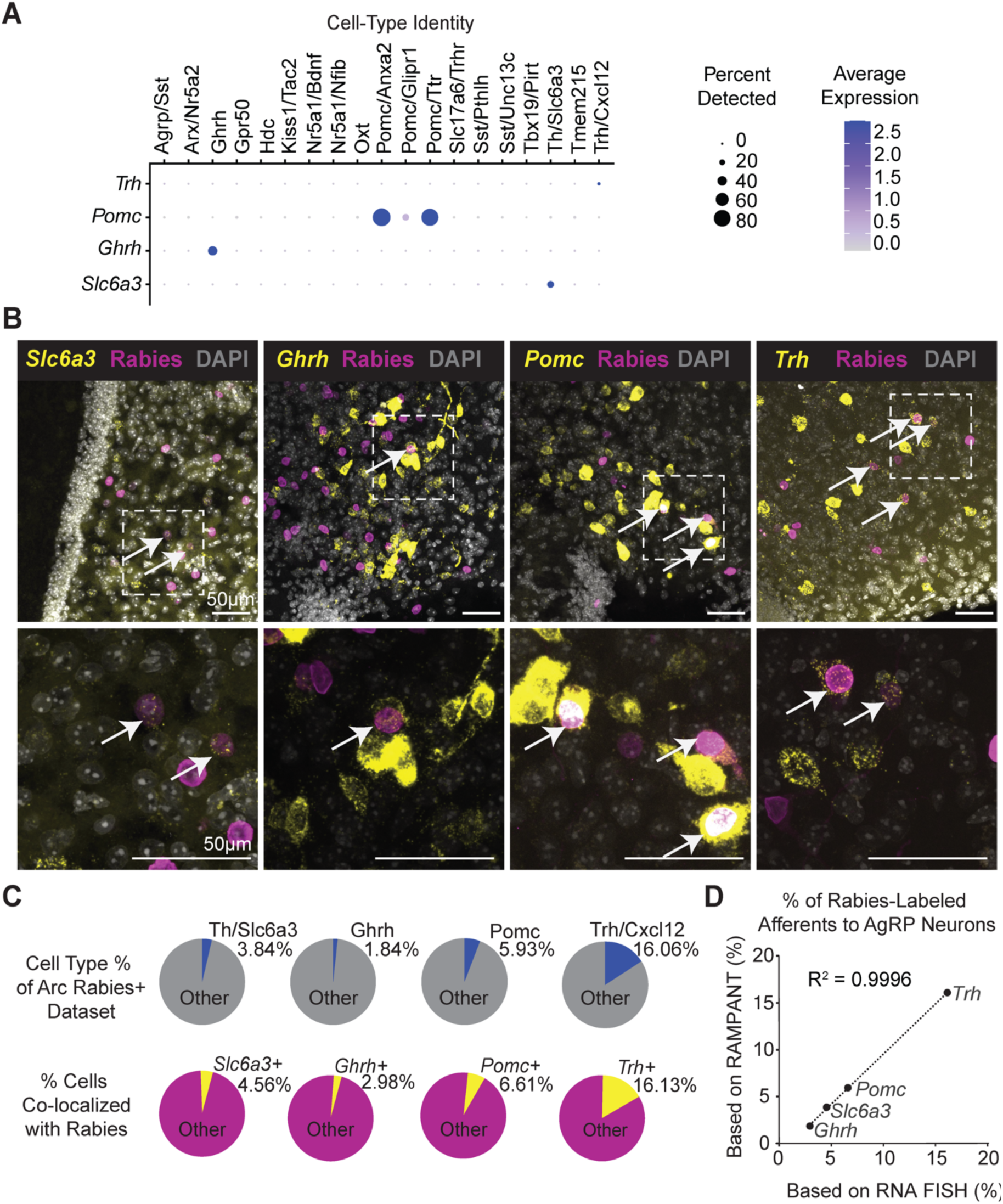
– Validation of RAMPANT Cell-Type Predictions by RNA FISH. **A**, Dot plot showing cluster-level expression of *Trh*, *Pomc*, *Ghrh*, and *Slc6a3*. **B,** Expression of *Slc6a3*, *Ghrh*, *Pomc*, and *Trh* transcripts (yellow) and rabies H2b-mCherry fluorescence (magenta) in Arc cells (n=3 mice). Top row imaged at 20x magnification; bottom row imaged at 63x magnification. **C,** (top) Pie charts of four cell clusters as percentages of the Arc RAMPANT dataset; (bottom) Pie charts of four cell-type markers as a percentage of cells after rabies labeling via AgRP neurons. **D,** Correlation between the percentages shown in the top and bottom pie charts in Figure 4C.

### RAMPANT Characterizes the Transcriptional Response to Metabolic Challenges

To evaluate if RAMPANT can detect metabolically regulated genes, we repeated our RAMPANT study on mice that had undergone one of three feeding conditions (fed, fasted, or post-fast refed) immediately prior to tissue collection (n=9-11 mice per feeding condition, Fig. 5A). The mice lost body weight with fasting and regained it with post-fast re-feeding, as expected (Fig. S5A). Comparing gene expression in rabies-infected AgRP neurons between fed and fasted mice revealed 134 fasting-sensitive genes (n=193 and 308 AgRP neurons from fed and fasted mice, respectively; Wilcoxon rank-sum test with Bonferroni correction; Fig. 5B-D). Of these genes, the majority (56%) were previously detected in a study comparing pooled AgRP neuron samples from fed and fasted mice, including genes upregulated by fasting: *e.g.*, *Mast4; Irs2; Npy; Vgf; Lepr;* and *Erg1* ^38^ (Fig. 5C). Additionally, our analysis revealed downregulation of many other genes with fasting (Fig. 5D), including *Ccdc85a*, *Rasgrp1*, *Camk2a*, and *Synpr*. These results confirm many genes previously reported to change in AgRP neurons with fasting ^30,38^.

**Figure 5.**
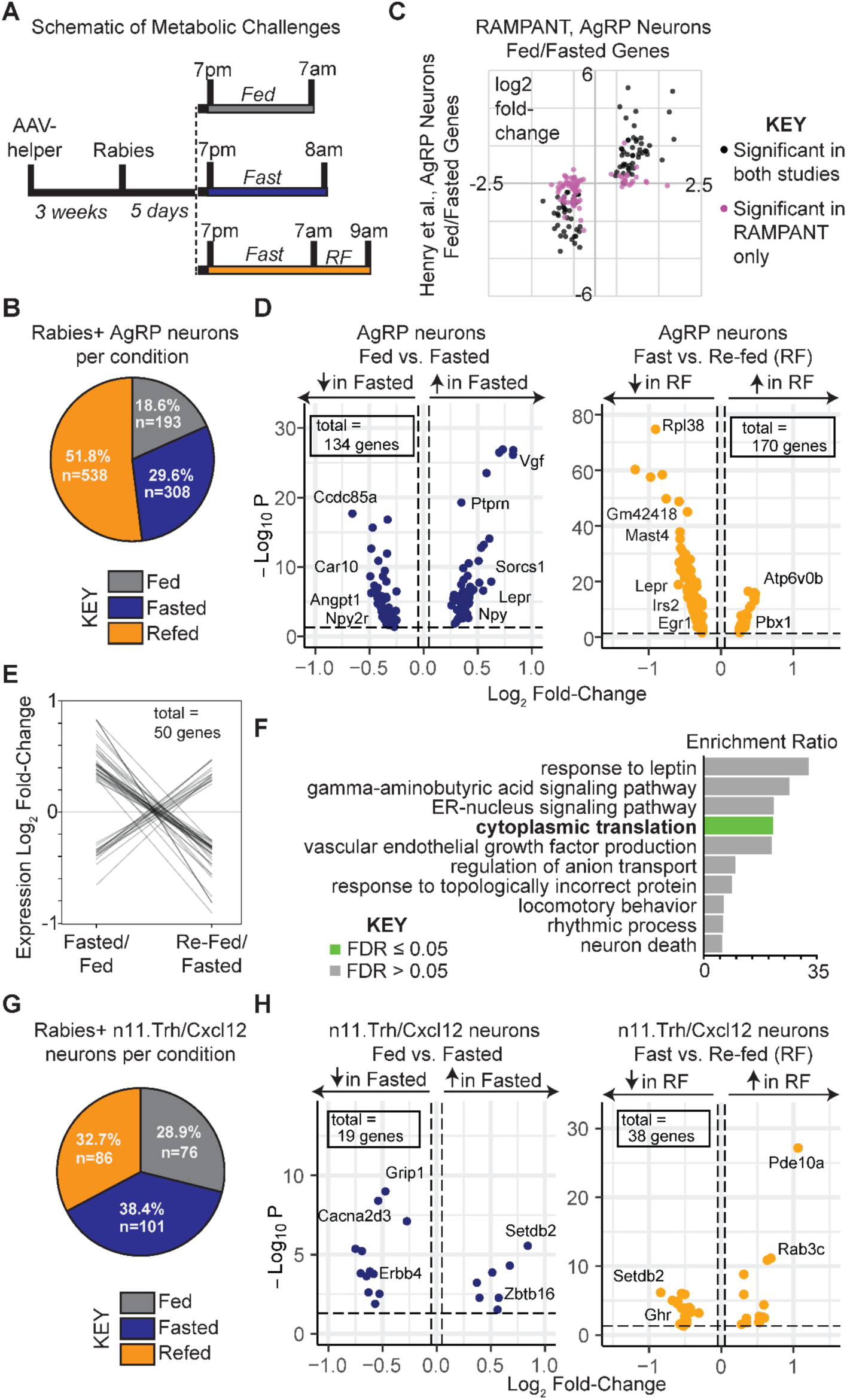
– RAMPANT Characterizes the Transcriptional Response To Metabolic Challenges. **A**, Timeline of virus injections and metabolic challenges; RF, re-feeding. **B,** Pie chart showing number of rabies-infected AgRP neurons per feeding condition (n=8 fed mice, n=8 fasted mice, n=10 refed mice). **C,** Comparison of log2 fold-change values in fasting-sensitive gene expression between the current study and ref. ^38^. **D,** (*left*) Volcano plot of genes differentially expressed between fed and fasted rabies-infected AgRP neurons (134 DE genes; Wilcoxon rank-sum test with Bonferroni correction); (*right*) volcano plot of genes differentially expressed between fasted and refed rabies-infected AgRP neurons (170 DE genes; Wilcoxon rank-sum, Bonferroni correction). **E,** Fasting and refeeding oppositely regulate expression levels of 50 genes in AgRP neurons; “fasted/fed” indicates log2 fold-change in expression from fed to fasted, whereas “refed/fasted” indicates log2 fold-change in expression from fasted to refed. **F,** Enriched gene ontology categories among the 50 genes oppositely regulated by fasting and refeeding in rabies-infected AgRP neurons. **G**, Pie chart showing distribution of rabies-infected n11.Trh/Cxcl12 neurons per feeding condition. **H**, Volcano plot of genes differentially expressed between fed and fasted rabies-infected n11.Trh/Cxcl12 neurons (left panel; 19 DE genes; Wilcoxon rank-sum with Bonferroni correction); volcano plot of genes differentially expressed between fasted and refed rabies-infected n11.Trh/Cxcl12 neurons (right panel; 38 DE genes; Wilcoxon rank-sum test with Bonferroni correction).

Comparing gene expression in rabies-infected AgRP neurons between fasted and refed mice revealed 170 feeding sensitive genes (Wilcoxon rank-sum, Bonferroni correction; Fig. 5D). This analysis showed that feeding downregulates the expression of many genes upregulated by fasting, such as *Mast4*, *Irs2*, *Lepr* and *Erg1* (Fig. 5D). Overall, feeding oppositely regulated 50 of the 134 fasting-sensitive genes to a statistically significant extent (Fig. 5E; Supplemental Table 1). Gene ontology analysis revealed enrichment for cytoplasmic translation among these 50 genes (*Rpl32*, *Rpl38*, *Rps21*, *Rps28*, *Rps2*9; Fig. 5F), suggesting that fasting and refeeding oppositely alter translational capacity in AgRP neurons. Our results thus demonstrate that RAMPANT can detect fasting– and feeding-sensitive genes and that feeding oppositely regulates over a third of fasting-sensitive genes in AgRP neurons.

We repeated our analysis on rabies-infected n11.Trh/Cxcl12 neurons, which were far less numerous than AgRP neurons in our dataset (n=263 n11.Trh/Cxcl12 neurons *vs*. 1,039 AgRP neurons; Fig. 5G). Our results revealed 19 differentially expressed genes between fed and fasted mice, as well as 38 differentially expressed genes between fasted and refed mice (Fig. 5H). We identified fewer differentially expressed genes in n11.Trh/Cxcl12 neurons than in AgRP neurons, likely due to the smaller sample size. Among the fasting-sensitive genes, we found an upregulation of some genes known to regulate body weight, including *Lepr*, *Tmem132d*, and *Erbb4* ^39–41^ (Fig. 5H). We also detected upregulation of *Rab3c* in refed mice when compared to fasted mice (Fig. 5H). Previous reports show that Rab3c modulates the IL6-STAT3 pathway ^42^, which is critical for glucose homeostasis in hepatic tissue and skeletal muscle ^43^. We also observed an increase in *Setdb2* expression with fasting, as well as a decrease in *Setdb2* expression after refeeding (Fig. 5H). While little work has been done studying the role of *Setdb2* in the hypothalamus, previous studies conducted in the liver have established *Setdb2* as a critical regulator of lipid metabolism through its role in the induction of *Insig2a* ^44^. This study also found an increase in *Setdb2* mRNA expression in fasted mice when compared to refed mice, suggesting this gene may play a regulatory role during fasting in multiple cell types ^44^. These results, along with their relatively high expression of genes encoding receptors for the satiety hormones GLP-1 and leptin (*Glp1r*, *Lepr*) suggest that n11.Trh/Cxcl12 neurons may have a role in energy balance (Fig. S4A).

### Trh Arcuate Neurons Synaptically Inhibit AgRP Neurons

AgRP neurons receive GABAergic input from *Lepr*+ and *Glp1r*+ Arc neurons ^3,18^, and our RAMPANT analysis predicts that n11.Trh/Cxcl12 neurons are a source of that input (Fig. S4A). To investigate this possibility, we used channelrhodopsin (ChR2)-assisted circuit mapping (CRACM) to determine whether Trh^Arc^ neurons form functional synapses onto AgRP neurons ^45,46^. After targeting ChR2 expression to the caudal Arc of Trh-Cre;Npy-hrGFP mice to selectively transduce Trh^Arc^ cells, we recorded post-synaptic currents from Npy-hrGFP Arc cells while photo-stimulating Trh^Arc^ axons in acute brain sections (Fig. 6A). Of note, the vast majority of Npy-hrGFP+ cells in the Arc are AgRP neurons ^47,48^. We did not observe any co-expression of ChR2 with Npy-hrGFP (data not shown) and further confirmed that AgRP neurons and Trh neurons are distinct populations using RNA FISH (Fig. S6D).

**Figure 6.**
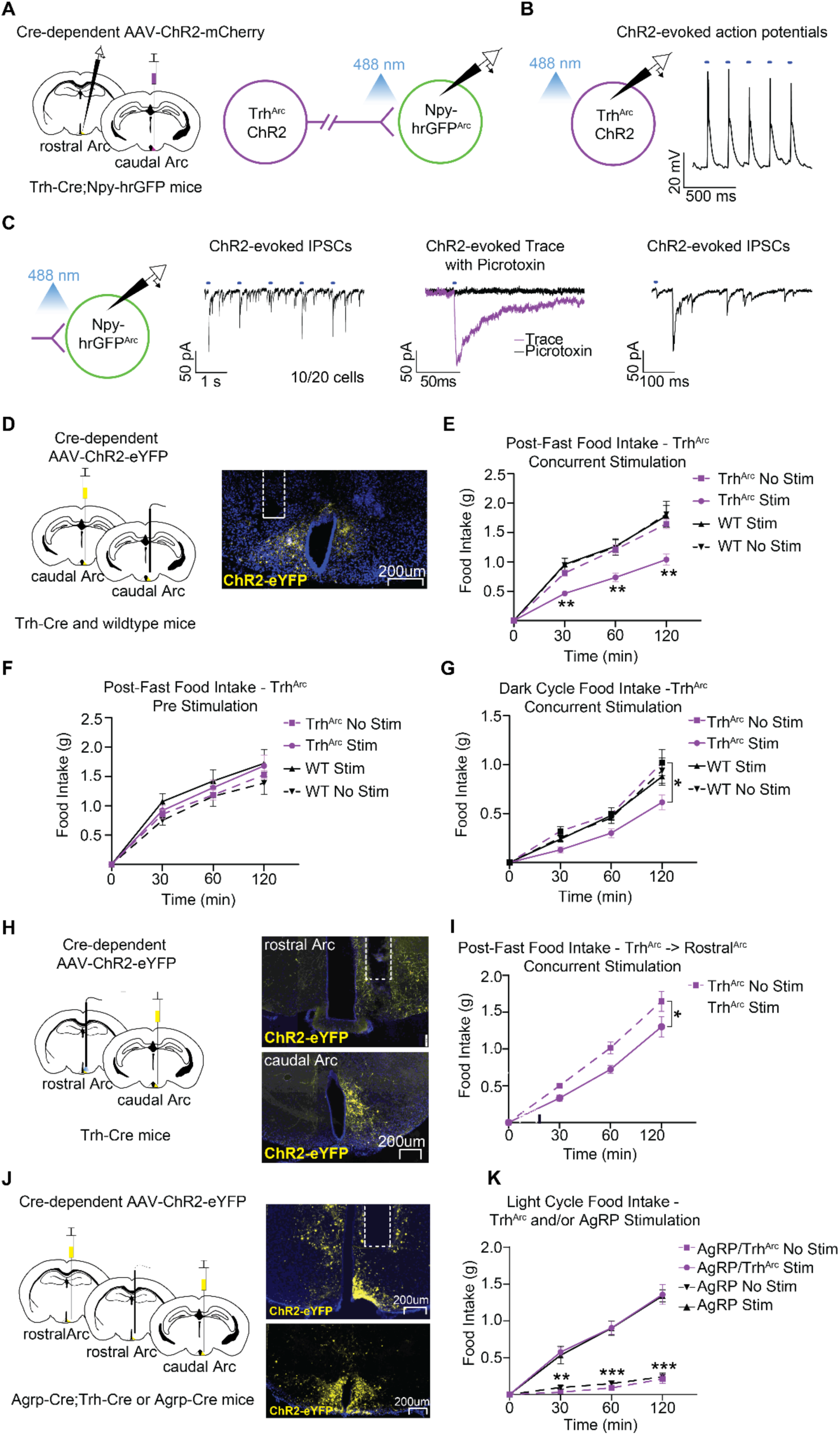
– Trh^Arc^ Neurons Suppress Appetite by Inhibiting AgRP Neurons. **A**, Left, schematic of unilateral viral delivery of Cre-dependent AAV-ChR2 to caudal Trh^Arc^ neurons (Trh^Arc^-ChR2) and electrophysiological recordings of rostral NPY-hrGFP neurons (∼AgRP neurons) in Trh-Cre;Npy-hrGFP mice. Right, schematic of channelrhodopsin-assisted circuit mapping (CRACM). **B,** Representative trace of ChR2 light-evoked action potentials in caudal Trh^Arc^-ChR2 neurons. **C,** Representative trace of ChR2 light-evoked IPSCs in NPY-hrGFP+ Arc neurons in the absence or presence of the GABA antagonist picrotoxin (n=20 cells from a total of 3 mice). **D,** Left, schematic of unilateral injection of Cre-dependent AAV-ChR2 to caudal Trh^Arc^ neurons and optical fiber implant over the caudal Arc in Trh-Cre mice. Right, representative image of Trh^Arc^ ChR2-eYFP expression and caudal fiber implant location. **E,** Average post-fast food intake during Trh^Arc^-ChR2 concurrent photostimulation (n=14 for Trh^Arc^, n=8 for wildtype, males and females, repeated-measures two-way ANOVA, time x condition: F (9, 120) = 5.97, P<0.0001, Tukey’s multiple comparisons). **F,** Average post-fast food intake during Trh^Arc^-ChR2 pre-photostimulation (n=12 for Trh^Arc^-ChR2, n=8 for wildtype, males and females, repeated-measures two-way ANOVA, time x condition: F (9, 108) = 0.67, P=0.74). **G,** Average dark cycle food intake during Trh^Arc^-ChR2 concurrent photostimulation (n=9 for Trh^Arc^-ChR2, n=7 for wildtype, males and females, repeated-measures two-way ANOVA, time x condition: F (9, 84) = 2.09, P=0.04). **H,** Left, schematic of unilateral delivery of Cre-dependent AAV-ChR2 to caudal Trh^Arc^ neurons and optical fiber implant over the rostral Arc in Trh-Cre mice. Right, representative images of Trh^Arc^ ChR2-eYFP expression and rostral Arc fiber implant location. **I,** Within-subject quantification of average post-fast food intake during rostral Arc Trh^Arc^-ChR2 concurrent photostimulation versus no stimulation (n=6, males and females, repeated-measures two-way ANOVA, Condition: F (1, 10) = 7.96, P=0.02). **J,** Left, schematic of unilateral delivery of Cre-dependent AAV-ChR2 to Agrp-Cre and Trh-Cre Arc neurons and optical fiber implant over the rostral Arc in Agrp-Cre;Trh-Cre mice. Right, representative images of AgRP ChR2-eYFP and Trh^Arc^ ChR2-eYFP expression in rostral and caudal Arc neurons, respectively, and fiber implant location in the rostral Arc. **K,** Average light cycle food intake during either AgRP-ChR2 or both AgRP-ChR2 and Trh^Arc^-ChR2 concurrent photostimulation (n=7, males and females, repeated-measures two-way ANOVA, time x condition: F (9, 72) = 31.59, P<0.0001, Tukey’s multiple comparisons). All error bars of **E-G**, **I**, and **K** represent standard error of the mean (SEM). *P<0.05 **P<0.01 ***P<0.001.

After demonstrating the ability of our approach to drive light-evoked action potentials in Trh^Arc^ neurons (Fig. 6B), we recorded from Arc Npy-hrGFP+ neurons and detected light-evoked inhibitory postsynaptic currents (IPSCs) in 50% Npy-hrGFP+ neurons tested (Fig. 6C). Importantly, these currents were blocked by the GABA A receptor antagonist, picrotoxin (Fig. 6C), indicating that Trh^Arc^ neurons release GABA onto AgRP neurons, consistent with their GABAergic molecular phenotype (e.g., *Slc32a1* expression) ^30^. Our results thus validate RAMPANT’s prediction that Trh^Arc^ neurons synapse on and inhibit at least a subset of AgRP neurons.

### Trh Arcuate Neurons Decrease Feeding in an AgRP Neuron-Dependent Manner

Activation of AgRP neurons promotes hyperphagia in calorically replete mice ^12,14^, while direct or indirect inhibition significantly reduces food intake ^14^. Since our CRACM studies demonstrate that Trh^Arc^ neurons inhibit AgRP neurons, we hypothesized that activating Trh^Arc^ neurons would reduce food intake in an AgRP neuron-dependent manner. To examine the ability of Trh^Arc^ neurons to diminish feeding, we targeted ChR2 expression to caudal Trh^Arc^ neurons and implanted optic fibers over the caudal Arc where Trh^Arc^ neurons predominantly reside (Fig. 6D). For controls, we injected Cre-negative littermate mice with the same Cre-dependent AAV and equipped mice with optic fibers targeting the caudal Arc. We then assessed food intake with and without concurrent photoactivation over a 2-hour period near the beginning of the light cycle following an overnight fast. Unilateral or bilateral Trh^Arc^ photoactivation during feeding significantly reduced post-fast food intake compared to the same mice without photostimulation and to controls with or without photostimulation (Fig. 6E). Our results show that activating Trh^Arc^ neurons can decrease feeding behavior.

AgRP neurons co-release NPY and AgRP, neuropeptides that induce a robust hyperphagic response when administered into the rodent hypothalamus ^49–51^. Moreover, activating AgRP neurons causes a sustained hunger drive which depends on NPY release ^52–54^. Since our RAMPANT data indicates that GABAergic Trh^Arc^ neurons contain transcripts for hormones and peptides (e.g., *Cartpt*, *Trh*), we investigated whether acutely activating these cells before food presentation causes a prolonged decrease in feeding. Accordingly, we primed Trh^Arc^ –ChR2 neurons with photoactivation in overnight fasted mice for one hour prior to food access, then stopped the photoactivation and allowed mice access to food for 2 hours. In contrast to our results with concurrent stimulation, photo-activating Trh^Arc^ –ChR2 neurons before food presentation failed to attenuate food intake in fasted mice, as all groups of mice exhibited comparable levels of food consumption (Fig. 6F). This suggests that the capacity of Trh^Arc^ neurons to suppress appetite is likely signaled through fast neurotransmitter communication as opposed to a delayed peptidergic mechanism.

While caloric deprivation between meals is typical for most animals, prolonged fasting leads to many physiological adaptations in mice including changes in hormone balance, body weight, metabolism, hepatic enzymes, cardiovascular parameters, body temperature, and toxicological responses ^55^. Mice under no caloric constraint consume most of their food at night, a significant portion of which occurs at the onset of the dark cycle ^56^. To examine the sufficiency of Trh^Arc^ neurons to reduce food intake in physiologically hungry mice that show circadian rhythmicity, food intake was measured with or without concurrent photoactivation in *ad libitum* fed mice for 2 hours after the onset of the dark cycle. As in the fast-refeed condition, Trh^Arc^ –ChR2 photoactivation reduced food intake during the first 2 hours of the dark cycle compared to the same mice without photostimulation, and to littermate wildtype controls as described above (Fig. 6G).

Since our CRACM studies showed that Trh^Arc^ neurons inhibit AgRP neurons through projections to the rostral Arc (see Fig. 6C), we investigated whether selectively activating these rostral Arc projections could also suppress feeding (Fig. 6H). Importantly, the AgRP neurons which innervate hunger-controlling brain regions and can drive feeding behavior are concentrated in the rostral half of the Arc ^57^. We found that photo-activating the rostral Arc axons of caudal Arc Trh^Arc^ –ChR2 neurons significantly reduced food intake after a fast (Fig. 6I), demonstrating the ability of these projections to decrease feeding in hungry animals.

If Trh^Arc^ neurons suppress feeding by inhibiting AgRP neurons in the rostral Arc, then stimulating AgRP neurons and Trh^Arc^ neurons together should prevent Trh^Arc^ neurons from reducing feeding. To test this prediction, we targeted ChR2 to Trh^Arc^ neurons and AgRP neurons in Trh-Cre;Agrp-Cre mice and placed an optical fiber over the rostral Arc (Fig. 6J). Simultaneous activation of AgRP neurons and Trh^Arc^ neurons dramatically increased food intake in sated mice, compared to mice without photostimulation. Importantly, this increase was almost identical to the hyperphagia seen in mice with only AgRP neuron photostimulation (Fig. 6K), indicating that AgRP neuron activity can override the satiating effects of sustained Trh^Arc^ neuron activity. Of note, our injection of ChR2 AAV into Trh-Cre;Agrp-Cre mice unavoidably also transduced a subset of Trh^DMH^ neurons with ChR2 (representative example in Fig. 6J). Whether Trh^DMH^ neurons contribute to feeding has not been reported. Nevertheless, our results indicate that activating Trh neurons in the Arc inhibit feeding through AgRP neurons.

Together, our results show that activating Trh^Arc^ neurons can suppress feeding through projections to the rostral Arc but in an AgRP neuron-dependent manner. Considering our finding that Trh^Arc^ neurons directly inhibit AgRP neurons, these results raise the possibility that Trh^Arc^ neurons decrease feeding mainly by inhibiting AgRP neurons. Supporting this notion, we found that photo-activating Trh^Arc^ neurons while chemogenetically inhibiting Npy^Arc^/AgRP neurons decreased post-fast food intake to a degree comparable to each manipulation alone (S7A-B).

### Trh Arcuate Neurons are Activated by Liraglutide and Contribute to Its Suppression of Appetite and Body Weight

Trh^Arc^ neurons express genes encoding receptors for two hormones whose critical targets in the brain remain unknown, glucagon-like peptide-1 (GLP-1 receptor gene, *Glp1r*; Fig. 7A, S4A) and leptin (leptin receptor gene, *Lepr*; Fig. S4A) ^30^. Liraglutide can diminish the response of AgRP neurons to food in hungry animals ^16^, potentially by activating inhibitory inputs to AgRP neurons ^3,16,17^. Since Trh^Arc^ neurons express *Glp1r* and inhibit AgRP neurons, we wondered whether they respond directly to liraglutide. To test this possibility, we imaged calcium activity in Trh^Arc^ neurons in acute brain sections, using the genetically encoded calcium indicator GCaMP6s (Fig. 7B). For each recording, slices were imaged in the presence of the synaptic blockers picrotoxin (25 uM), AP5 (20 uM) and NBQX (10 uM) for the entire recording session. After a 5-minute baseline period, during which slices were imaged in the presence of artificial cerebrospinal fluid (aCSF), either liraglutide or a saline vehicle was added to the bath for 10 minutes. Liraglutide (100 nM) application led to a steady ramp up of calcium activity over the wash-on phase compared to saline (Fig. 7C-D). Importantly, KCl was added to each slice at the conclusion of the recordings as a positive control to demonstrate cell responsiveness, even in those that failed to respond to liraglutide (Fig. 7C). To account for differences in cell responsiveness across tissue slices, we also calculated the percentage of responding cells per slice. Significantly more cells per slice responded to liraglutide than to saline (Fig. 7D). Of note, the percentage of liraglutide-responsive Trh^Arc^ neurons was consistent with the percentage of *Glp1r*-expressing Trh^Arc^ neurons we found by RNA FISH (see Fig. 7A).

**Figure 7.**
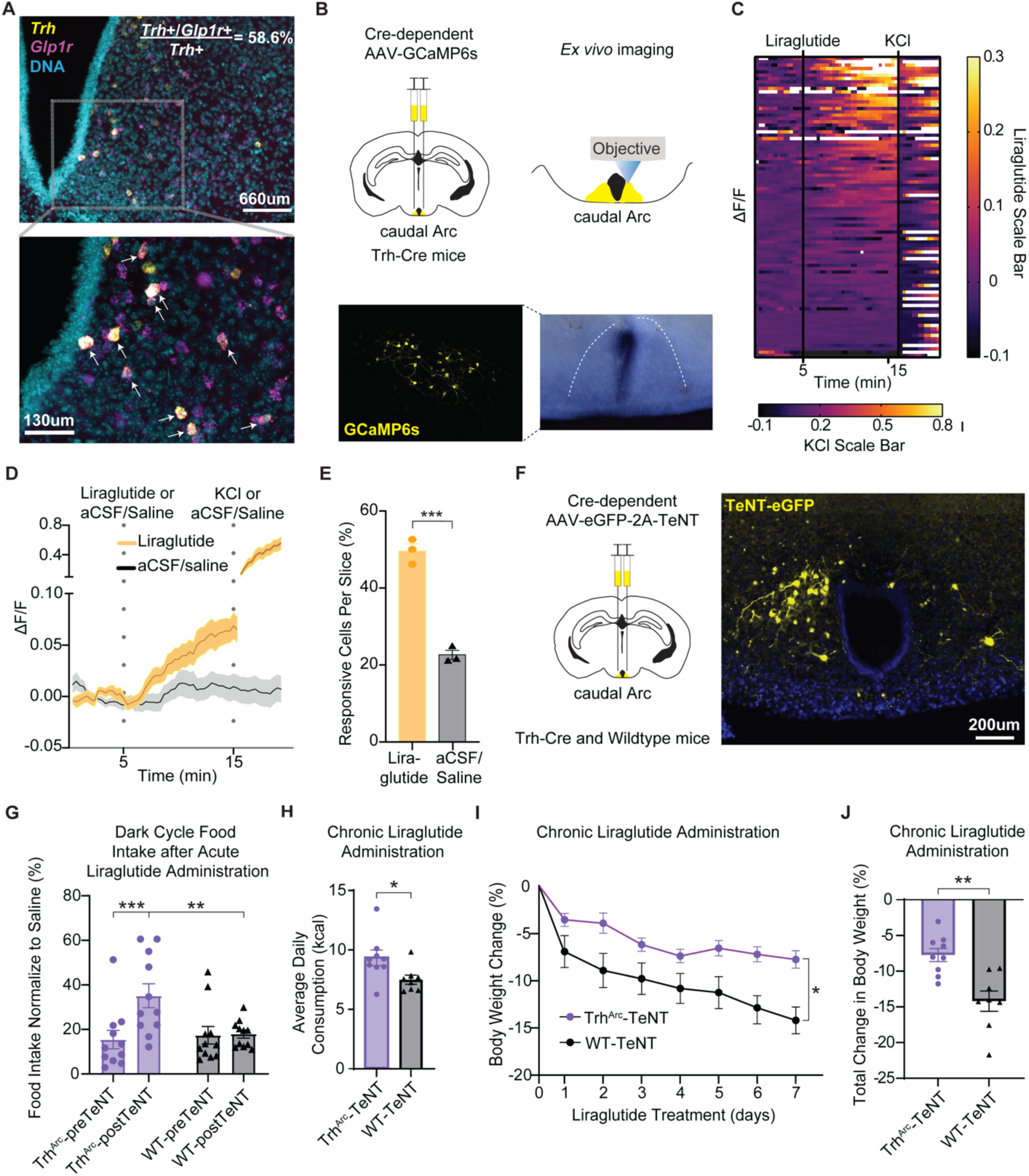
– Trh^Arc^ Neurons Respond to GLP-1R Agonists and Contribute to Their Suppression of Feeding and Body Weight. **A**, Representative image of *Glp1r* and *Trh* RNA FISH in the Arc, with arrows indicating co-expressing cells in the lower panel (percentage based on n=901 cells from 4 mice). **B,** Top, brain schematic of bilateral viral delivery of Cre-dependent GCaMP6s to caudal Trh^Arc^ neurons in Trh-Cre mice. Bottom, representative image of GCaMP6s expression in caudal Trh^Arc^ neurons. **C,** Heatmap of individual Trh^Arc^ neuron ΔF/F responses to liraglutide (100 nM) application and KCl (10 mM). Liraglutide scale bar for minutes 0 to 15, KCl scale bar for minutes 15 to 20 (n=90 cells/3 slices/3 mice, males and females). **D,** Averaged individual traces of Trh^Arc^ neuron ΔF/F responses to liraglutide followed by KCl vs. saline application (n=90 cells/3 slices/3 mice for liraglutide, n=64 cells/3 slices/3 mice for saline). **E.** Percentage of responsive cells per slice (n=3 for liraglutide, n=3 for saline, unpaired t-test (two-tailed), p=0.0003). **F,** Left, brain schematic of bilateral viral delivery of Cre-dependent AAV-eGFP-2a-TeNT to caudal Arc neurons in Trh-Cre mice. Right, representative image of Trh^Arc^-eGFP-2a-TeNT expression in caudal Arc neurons. **G**, Overnight food intake following acute liraglutide injection, calculated as the percentage of food intake following saline injection at the corresponding time-point, beeline (pre-TeNT) vs post-TeNT (n=11 for Trh^Arc^-TeNT, n=11 for WT-TeNT, males and females, RM two-way ANOVA, Time x Condition: F (1, 10) = 6.98, p=0.02, Tukey’s multiple comparisons). **H,** Average body weight change over 1 week of daily liraglutide administration in Trh^Arc^-TeNT and wildtype (WT) littermates bilaterally injected with Cre-inducible AAV-eGFP-2a-TeNT (n=9 for Trh^Arc^-TeNT, n=8 for WT-TeNT, males and females, repeated-measures two-way ANOVA, time x condition: F (6, 90) = 2.608, p=0.02). **I,** Total percentage body weight change over 1 week of daily liraglutide administration in Trh^Arc^-TeNT and wildtype (WT) littermates bilaterally injected with Cre-inducible AAV-EGFP-2a-TeNT (n=9 for Trh^Arc^-TeNT, n=8 for WT-TeNT, males and females, unpaired two-tailed t-test, t(15)=3.9, p<0.002). **J,** Average daily kcal consumption over 1 week of daily liraglutide administration in Trh^Arc^-TeNT and wildtype (WT) littermates bilaterally injected with Cre-inducible AAV-eGFP-2a-TeNT (n=9 for Trh^Arc^-TeNT, n=8 for WT-TeNT, males and females, unpaired two-tailed t-test, t(15)=2.4, p<0.05). All error bars and the shaded regions in panel D represent standard error of the mean (SEM). *p<0.05, **p<0.01, ***p<0.001, ****p<0.0001.

Since Trh^Arc^ neurons also express the gene encoding leptin receptor (Fig. S4A), we repeated our calcium imaging studies while administering leptin or its vehicle to the brain slice. However, in contrast to our results with liraglutide, leptin failed to significantly affect calcium activity in Trh^Arc^ neurons (data not shown). Importantly, our results do not rule out a delayed effect of leptin on these neurons. While leptin can directly alter the activity of some Arc neurons ^58,59^, it controls appetite largely through transcriptional changes ^60^, which presumably occur on a longer timescale than our imaging experiments.

GLP-1R agonists (GLP-1RA), such as liraglutide, can dramatically reduce body weight and food intake in rodents and humans ^61,62^. To determine whether Trh^Arc^ neurons are necessary for GLP-1RA-induced weight loss and satiety, we genetically targeted expression of tetanus toxin (TeNT) to caudal Trh^Arc^ neurons (Fig. 7F). TeNT irreversibly inhibits synaptic release by cleaving the synaptic vesicle protein synaptobrevin ^63^. We chose this approach over cell ablation to avoid collateral damage from gliosis and other immune responses to cell death. For controls, we injected wildtype littermates with the same virus in the caudal Arc.

To explore whether Trh^Arc^ neurons are necessary for liraglutide to acutely control feeding, we compare the effects of liraglutide on feeding prior to TeNT surgery (baseline) and at the conclusion of the body weight study, approximately 11 weeks after TeNT surgery. For each trial, we removed food from the home cage 2.5 hours prior to the onset of the dark cycle, injected the mice with liraglutide or vehicle 0.5 hours prior to the onset of the dark cycle, and measured food intake over the course of the dark cycle with a FED3 system ^64^. To account for changes in body weight affecting food intake ^65^, we calculated food intake after liraglutide treatment as a percentage of food intake following vehicle injection for the corresponding time point (i.e., baseline or 11 weeks post-transduction). As expected, acute administration of liraglutide potently reduced food intake in both groups of mice before TeNT transduction (pre-TeNT; Fig. 7G, S7C). However, while liraglutide suppressed food intake to a similar degree in wildtype-TeNT controls 11 weeks later, this suppression was significantly less in Trh^Arc^-TeNT mice (post-TeNT; Fig. 7G, S7C). These results demonstrate that Trh^Arc^ neuron signaling is necessary for the full extent of liraglutide’s acute anorectic effect.

To determine whether Trh^Arc^ neurons mediate liraglutide’s control of body weight, we administered liraglutide daily over the course of one week to mice on a high-fat diet. In control mice daily liraglutide treatment led to rapid and sustained body weight loss, at least in part due to decreased food consumption (Fig. 7H-J, S7D-E). However, silencing Trh^Arc^ neurons significantly blunted the effects of liraglutide on body weight, by about 50%, and food intake (Fig. 7H-J, S7D-E), suggesting these cells play a role in the effects of GLP-1RAs on energy balance. However, this manipulation only partially blocked liraglutide’s suppression of body weight and appetite, indicating that other cells must also be necessary for liraglutide’s full effects. Overall, our results demonstrate that (1) liraglutide can rapidly activate Trh^Arc^ neurons *ex vivo*, independent of fast-acting neurotransmission, and that (2) Trh^Arc^ neuron signaling contributes to both the chronic and acute anorectic effects of liraglutide.

Trh^Arc^ Neurons Are Activated by Feeding, Signal Satiety, and Receive Mostly Local Input Since Trh^Arc^ neurons contribute to liraglutide-induced satiety and weight loss, we wondered whether these neurons are generally necessary for energy balance. Indeed, significantly more *Trh*+/*Glp1r*+ Arc neurons expressed the immediate early gene and cell activity marker, *Fos*, after a post-fast meal than during fasting, indicating that Trh^Arc^ neurons are activated by feeding (Fig. 8A-B). We therefore investigated whether Trh^Arc^ neurons are necessary for satiety and body weight control. We did this by synaptically silencing Trh^Arc^ neurons with TeNT and tracking total body weight and food intake on a weekly basis at baseline and for 8 weeks, starting 3 weeks after the surgery date to allow for efficient viral transduction. We found that silencing Trh^Arc^ neurons significantly escalated body weight gain in mice on a standard chow diet (Fig. 8C, S7F), which coincided with elevated food intake (Fig. 8D, S7G). These results indicate that Trh^Arc^ neurons signal satiety and help to limit body weight gain *in vivo*.

**Figure 8.**
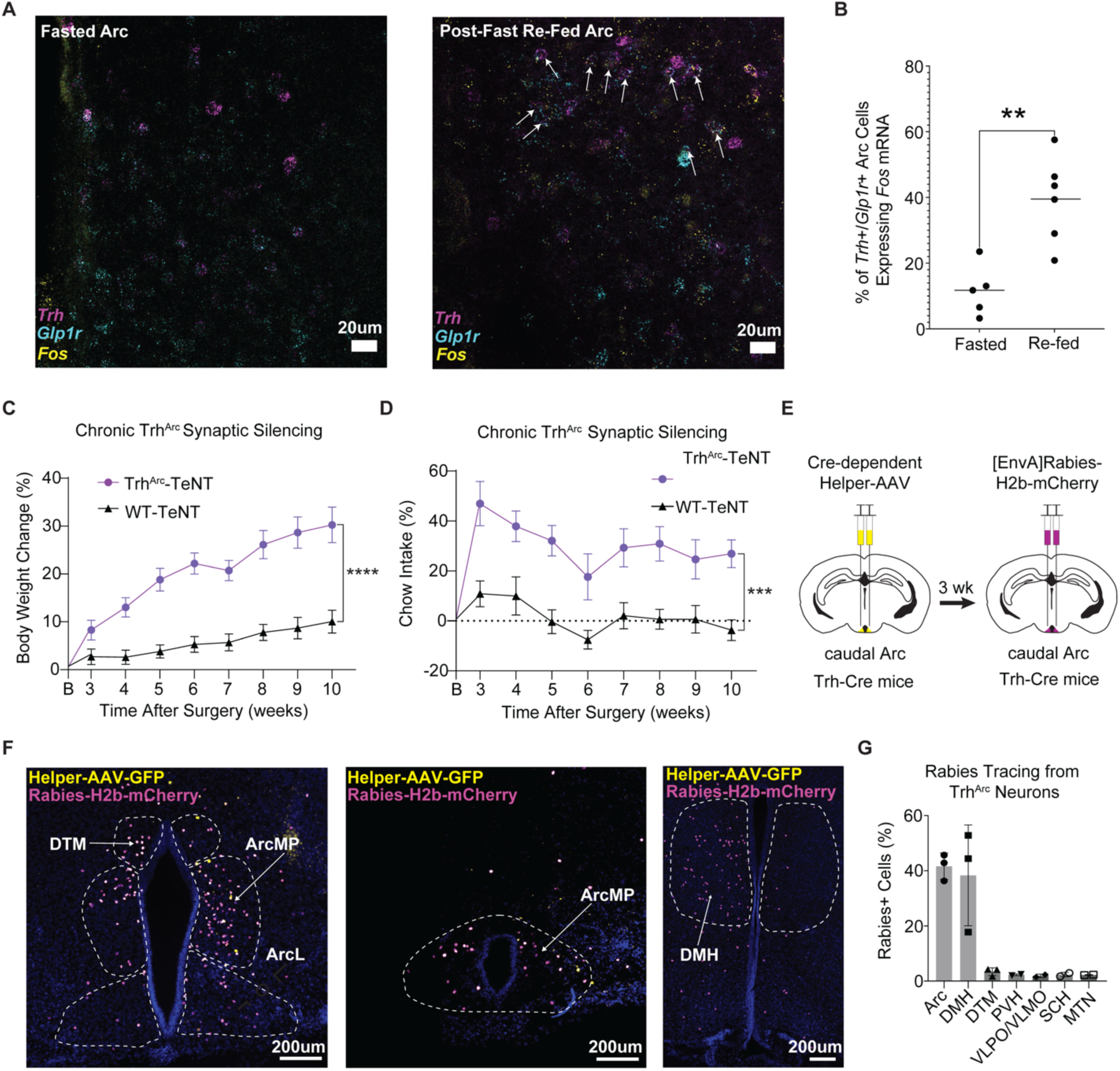
– Trh^Arc^ Neurons Are Activated by Feeding, Signal Satiety, and Receive Mostly Local Input. **A**, Representative images of Trh, Glp1r, and Fos RNA FISH colocalization in the Arc after overnight fasting, or overnight fasting plus 2 hr of re-feeding. **B,** The *Fos* mRNA+ percentage of *Trh*+/*Glp1r*+ Arc cells after fasting or post-fast re-feeding. **C,** Average body weight change over time normalized to baseline (“B”) body weight in Trh^Arc^-TeNT and wildtype (WT) littermates bilaterally injected with Cre-inducible AAV-eGFP-2a-TeNT (n=17 for Trh^Arc^-TeNT, n=17 for WT-TeNT, males and females, repeated-measures (RM) two-way ANOVA, time x condition: F (7, 70) = 3.00, p<0.0001). **D**, Average daily chow intake over time in Trh^Arc^-TeNT and wildtype (WT) littermates bilaterally injected with Cre-inducible AAV-eGFP-2a-TeNT (n=17 for Trh^Arc^-TeNT, n=17 for WT-TeNT, males and females, repeated-measures two-way ANOVA, time x condition: F (8, 80) = 7.73, p<0.0001). **E,** Schematic of bilateral viral delivery of Cre-dependent TVA helper AAV and rabies virus to caudal Trh^Arc^ neurons in Trh-Cre mice. **F,** Representative images of TVA helper AAV (yellow) and rabies virus (magenta) in the caudal Arc and rabies virus in the DMH. **G,** Percentage of total rabies+ cells found in each region per mouse (n=3 mice, 1 slice per mouse). Arc, Arcuate; DMH, Dorsomedial Hypothalamus; DTM, Dorsal Tuberomammillary; PVH, Paraventricular Hypothalamus; VLPO/VMPO, Ventrolateral / Ventromedial Preoptic Nucleus; SCH, Suprachiasmatic Nucleus; MTN, Midline Group of the Dorsal Thalamus. **p<0.01, ***p<0.001, ****p<0.0001.

While most of the body’s GLP1 comes from L-cells of the small intestine, it is also secreted by neurons in the medullary nucleus of the solitary tract (NTS) which control feeding ^66–68^. To determine whether Trh^Arc^ neurons receive input from GLP1+ NTS neurons, we mapped synaptic afferents to Trh^Arc^ neurons using monosynaptic retrograde rabies tracing. We restricted rabies infection to Trh^Arc^ neurons and their presynaptic partners by injecting a Cre-dependent helper virus expressing the TVA receptor and oG into the caudal Arc of Trh-Cre mice and one month later injecting the same region with [EnvA]rabies-H2b-mCherry (Fig. 8E). Our results indicated that a large majority of H2b-mCherry+ cells were in the vicinity of the Trh^Arc^ neurons (Fig. 8F-G). Quantifying expression levels revealed the largest percentage of rabies-infected cells were in the Arc and DMH, with far smaller subsets in nearby hypothalamic regions, including the dorsal tuberomammillary (DTM), PVH, ventrolateral/ventromedial preoptic nucleus (VLPO/VMPO), suprachiasmatic nucleus (SCH), and midline group of the dorsal thalamus (MTN) (Fig 7N). Notably, we failed to detect rabies-labeled cells in the medulla, suggesting Trh^Arc^ neurons receive the bulk of their synaptic input from other mediobasal hypothalamic neurons. While it remains possible that Trh^Arc^ neurons receive extrasynaptic or polysynaptic input from medullary GLP1 neurons, it is also conceivable that the Trh^Arc^ neurons and their afferents integrate information from circulating signals via the third ventricle and median eminence.

## DISCUSSION

We molecularly classified afferents to AgRP neurons in the mediobasal hypothalamus using RAMPANT, a novel method combining rabies-based connectomics with single-cell transcriptomics. Among the afferents was a previously uncharacterized subtype of inhibitory neurons in the caudal Arc marked by expression of *Trh*, *Glp1r*, and *Lepr*. We confirmed that Trh^Arc^ neurons provide direct GABAergic input to AgRP neurons and found that activating Trh^Arc^ neurons decreased feeding in an AgRP neuron-dependent manner. Finally, we demonstrated that Trh^Arc^ neurons (1) can directly respond to liraglutide, (2) contribute to liraglutide’s satiating effects and weight loss (but not that of exogenous leptin), and (3) limit feeding and body weight gain. By identifying this liraglutide-sensing Trh^Arc^→ AgRP neuron satiety circuit, our studies provide a potential mechanism for the effectiveness of weight loss-inducing GLP-1R agonists.

Viruses other than rabies can be used to trans-synaptically label afferent neurons for molecular classification. For instance, previous studies have used pseudorabies virus (PRV) to label afferents to *Crh*-expressing neurons of the paraventricular hypothalamus, with methods called Connect-seq (dissociated cells) ^69^ and nuConnect-seq (cell nuclei) ^70^. Like RAMPANT, nuConnect-seq labels afferents to a genetically defined neuron population with a retrograde transsynaptic virus, profiles the labeled cells by single-nucleus RNA-seq and identifies them by transferring cell-type labels from a reference dataset of uninfected cells. However, nuConnect-seq and RAMPANT differ in fundamental ways. First, PRV can spread poly-synaptically, so the PRV-infected neurons in nuConnect-seq may be multiple synapses away from the starter neurons. For instance, the *Npy*+ Arc neurons previously identified by Connect-seq as afferents to *Crh*+ PVH neurons ^69^ may in fact be AgRP neurons which are two synapses upstream of *Crh*+ PVH neurons ^71^. In contrast, G-deficient mutants of rabies virus (ΔG rabies), such as the one we used in RAMPANT, are monosynaptically restricted, since they can only spread from starter cells expressing rabies glycoprotein ^19–21^. Therefore, unlike nuConnect-seq, RAMPANT profiles only starter cells and their primary afferents. Another key difference is that nuConnect-Seq captures more non-neuronal cells (i.e., glia and endothelial cells) than neurons. This may be due to the spread of PRV into neighboring, unconnected cells ^70^, raising the possibility that PRV could similarly infect bystander neurons and so confound interpretation of the results. In contrast, our RAMPANT study of AgRP neurons afferents detected only neurons, consistent with previous studies showing that G-deficient rabies spreads specifically between connected neurons ^20,72^.

Determining synaptic connectivity in neural networks, including synaptic inputs to AgRP neurons, is a fundamental challenge in neuroscience. The current standard for identifying connected neurons is to infect them with a modified rabies virus (e.g., SADΔG-B19) which spreads monosynaptically, labeling only the initially infected neurons and their presynaptic partners ^19–21^. While this method reveals the location and anatomy of synaptic afferents, it provides little insight into their molecular identities. Recent studies have profiled gene expression in rabies-infected cells by single-cell RNA-sequencing (scRNA-seq). Their results show that, while rabies infection alters the expression of some genes, infected cells can still be classified by comparing their gene expression profiles to those of uninfected cells ^24,29,73–75^. Our use of RAMPANT confirms that rabies-infected cells can be molecularly identified and extends these previous findings, demonstrating that most AgRP neuron marker genes are unaffected by rabies 5 days after infection and that their molecular response to fasting and feeding remains largely intact. Importantly, this method can be applied to any neuron of interest in order to molecularly classify its synaptic afferents in a high-throughput manner.

Our RAMPANT results predict 14 molecular subtypes of Arc neurons that synapse on AgRP neurons. The extensive interconnectivity suggested by our results is consistent with previous anatomical studies of the Arc ^76^. For instance, roughly half of all synapses in the Arc are thought to originate from other Arc neurons ^77^. Prominent among the known afferents we detected were KNDy neurons, which release glutamate onto metabotropic receptors on AgRP neurons, thereby inhibiting them ^33^. In our RAMPANT analysis of AgRP neurons and their afferents, n20.Kiss1/Tac2 neurons were the third most abundant Arc neuron subtype.

Another source of synaptic input to AgRP neurons may be dopaminergic Arc neurons. Specifically, dopaminergic axons arborize and form terminal-like structures around AgRP neuron bodies ^34^. However, optogenetically activating axons from Arc tyrosine hydroxylase (TH)+ neurons failed to alter AgRP neuron membrane potential in acute brain sections ^34^. Our RAMPANT analysis identifies tuberoinfundibular dopaminergic (TIDA) neurons (n08.Th/Slc6a3 neurons) as a potential source of local dopaminergic input to AgRP neurons. However, while the rabies transmission from AgRP neurons to n08.Th/Slc6a3 neurons we observed is consistent with a synaptic connection between these populations, further investigation is needed to validate that connection and determine its functionality.

Finally, Arc neurons that express dopamine receptor D1 (*Drd1*+) neurons also provide glutamatergic and GABAergic input to AgRP neurons ^35^. Among the *Drd1*+ subtypes in our RAMPANT dataset are POMC neurons (n15.Pomc/Anxa2, n21.Pomc/Glipr1), inhibitory neuron subtypes (*e.g.*, n24.Sst/Pthlh, n27.Tbx19), and excitatory neuron subtypes (*e.g.*, n32.Slc17a6/Trhr, n29.Nr5a1/Bdnf; Fig. S4). Our results thus predict specific subtypes of *Drd1*+ neurons which could transduce dopamine signaling for AgRP neurons.

The Arc contains abundant GABAergic neurons, many of which increase feeding when activated, including AgRP neurons ^12,14^, somatostatin/*Sst*+ Arc neurons ^30,78^, and tyrosine hydroxylase/*Th*+ Arc neurons ^34,79^. Chronically activating non-AgRP GABAergic Arc neurons causes hyperphagia and obesity ^80^, indicating an orexigenic role for GABAergic Arc neurons in general. For instance, Arc neurons expressing the prepronociceptin gene (*Pnoc*), the vast majority of which are GABAergic (*Slc32a1*/vGAT+) and distinct from AgRP, POMC, and TH Arc neurons, are activated by high-fat diet, can increase food intake upon activation, and are necessary to limit feeding and body weight gain on a high-fat diet ^81^. To our knowledge, Trh^Arc^ neurons are the first GABAergic Arc neuron found to have the opposite effect and suppress feeding when activated.

While some studies have implicated the Arc in liraglutide’s suppression of appetite ^3^, others indicate that Arc GLP-1Rs do not control appetite. For instance, Sandoval and colleagues reported that injecting glucagon-like peptide 1 (GLP-1) into the Arc of rats did not significantly affect their feeding ^82^. However, this study targeted GLP-1 ventrally in the caudal Arc, near the bottom of the third ventricle ^82^. Since the Trh^Arc^ neurons targeted in our study predominantly reside at the top of the third ventricle, it is possible that they were not affected by the GLP-1 injection. Our study demonstrates that signaling from Trh^Arc^ neurons participates in liraglutide’s satiating effects but does not rule out the possibility that liraglutide acts upstream of Trh^Arc^ neurons in the circuit. For instance, the liraglutide-induced calcium transients we observed in Trh^Arc^ neurons *ex vivo* may have been due to neuropeptidergic signaling from liraglutide-sensing neurons to Trh^Arc^ neurons, signaling which would have been spared by the synaptic blockers we used. Consistent with this, gene knockout studies indicate that *Glp1r* expression by glutamatergic (*Slc17a6*+) neurons but not by GABAergic (*Slc32a1*+) neurons is required for liraglutide to suppress appetite ^61^. Further investigation is needed to determine whether Trh^Arc^ neurons can directly sense liraglutide *in vivo* and whether this is necessary for the resulting satiety.

To investigate neurons that co-express *Lepr*, *Glp1r,* and *Slc32a1,* such as the Trh^Arc^ neurons described here, a recent study used conditional knock-out and rescue strategies to better define the role of these neurons in body weight maintenance ^5^. The authors found that deleting *Lepr* expression from *Glp1r*+ cells stimulated weight gain through overeating. Expressing *Lepr* in GABAergic (*Slc32a1*+) neurons in otherwise *Lepr*-null mice almost completely rescued them from obesity and hyperphagia, akin to complementary studies deleting *Lepr* from GABAergic neurons ^83^. However, no such rescue occurred if GABAergic *Glp1r*+ neurons were excluded from the *Lepr* reactivation, indicating that leptin acts on GABAergic *Glp1r*+ neurons to control feeding and body weight. Importantly, a similar knockout and conditional reactivation approach showed that expressing *Glp1r* only in GABAergic *Lepr*+ cells was sufficient for liraglutide to suppress appetite ^5^. The study concluded that the likely site of interaction of liraglutide and leptin was the DMH given the abundance of leptin-sensing *Glp1r*+ neurons there ^5^. However, the study does not rule out a role for Trh^Arc^ neurons characterized in the present study, since they also co-express *Slc32a1*, *Lepr*, and *Glp1r*. Indeed, these Trh^Arc^ neurons are the only Arc neurons to express the gene *Bnc2* ^30^ and so, as reported in a recent preprint, may be a direct target for leptin in its control of appetite and body weight ^84^.

Other hypothalamic neurons may also couple GLP-1R signaling to inhibition of AgRP neurons. For instance, a recent study found that *Glp1r*+ neurons in the DMH (Glp1r^DMH^ neurons) are activated by GLP-1R agonists to suppress feeding and body weight, potentially through their synaptic inhibition of AgRP neurons ^28^. Our RAMPANT results validate these previous findings by showing that Glp1r^DMH^ neurons are afferents to AgRP neurons (Fig. S3G). However, the Glp1r^DMH^ neurons in our study also expressed the leptin receptor gene, *Lepr*, and so may correspond to a previously identified population of DMH inhibitory neurons which co-express *Glp1r* and *Lepr* ^85^. *Lepr*+ DMH neurons inhibit AgRP neurons in response to external food cues to reinforce food-seeking behavior ^18,86^. However, in contrast to *Lepr*+ DMH neurons^86^, the Trh^Arc^ neurons characterized in our study received little if any synaptic input from the lateral hypothalamus (Fig. 7L). This raises the possibility that the Glp1r^DMH^ neurons and Trh^Arc^ neurons convey different information to AgRP neurons. For instance, while the Glp1r^DMH^ neurons relay pre-ingestive sensory cues to AgRP neurons ^28^, the Trh^Arc^ neurons may instead transmit post-absorptive metabolic cues, such as leptin ^84^, consistent with our observation that their synaptic input largely comes from the Arc and DMH (Fig. 7L). Further investigation is needed to understand the physiological role of Trh^Arc^ neurons and how it differs from that of Glp1r^DMH^ neurons. Along with other *Glp1r*+ neural cell populations (e.g., ^8,61,87,88^), we propose that Trh^Arc^ neurons are part of a distributed network of neural cells through which GLP-1 signaling suppresses feeding.

## METHODS

All animal care and experimental procedures were approved in advance by the University of Virginia Institutional Animal Care and Use Committee (RAMPANT and RNA FISH experiments) and by the US National Institutes of Health Animal Care and Use Committee (all other experiments). Mice were housed with a 12 hr light/dark cycle and provided *ad libitum* access to food (standard chow, Envigo 7017 NIH-31, or 20 mg grain pellets, TestDiet 5TUM) and water unless otherwise noted. All experiments were carried out in adult (>8 weeks) mice that were group-housed until experiments began. Some measurements were carried out in the same mouse across conditions (see individual methods sections for further details).

### Genotypes

C57BL/6J, Agrp-IRES-Cre (“Agrp-Cre”, Jackson Laboratories, JAX, stock # 012899), Trh-IRES-Cre (“Trh-Cre”, gift from Bradford B. Lowell), Npy-hrGFP (JAX, stock # 006417) and Npy-FlpO (JAX, stock # 030211) mice were used. Agrp-Cre mice have an internal ribosome entry site (IRES)-Cre inserted after the stop codon of the *Agrp* gene on chromosome 8 ^89^. Trh-Cre mice have an internal ribosome entry site (IRES)-Cre inserted after the stop codon of the *Trh* gene on chromosome 6 ^23^. Npy-hrGFP mice express humanized Renilla Green Fluorescent Protein (hrGFP) under control of the mouse *Npy* gene promoter ^48^. Npy-IRES2-FlpO-D knock-in mice (“Npy-Flp”, Jackson Laboratories, JAX stock # 030211) have an optimized FLP recombinase targeted to *Npy-*expressing cells ^90^. For the electrophysiology experiments, Trh-Cre mice were crossed with Npy-hrGFP mice to produce Trh-Cre;Npy-hrGFP mice. To inhibit Agrp neurons during Trh neuron activation, Npy-Flp mice were crossed with Trh-Cre mice to produce Npy-Flp;Trh-Cre mice. For concurrent photostimulation experiments, Agrp-Cre mice were crossed with Trh-Cre mice to produce Agrp-Cre;Trh-Cre mice.

### Drugs

Leptin (R&D Systems, catalog # 498-OB), Liraglutide (Selleckchem, catalog # S8256), KCl (Sigma Aldrich, catalog # 7447-40-7), and NBQX (Abcam, catalog # ab120046), and DL-AP5 (Abcam, catalog # ab120004) was dissolved in saline at stock concentrations and stored at –20°C until use. Picrotoxin (Abcam, catalog # ab120315) was dissolved in DMSO and stored at room temperature until use. For *in vivo* experiments, on the day of testing, the stock solution was thawed, diluted with saline, and delivered at a volume of 10 ml/kg at the following doses: Leptin (4.0 mg/kg) ^30^, Liraglutide (0.2 mg/kg) ^91^, and CNO (3.0 mg/kg). For *in vitro* slice experiments, drugs were diluted in 1X artificial cerebrospinal fluid (aCSF) at the following concentrations: leptin (100 nM) ^92^, liraglutide (100 nM) ^17^, KCl (10 mM), picrotoxin (25 uM), AP5 (20 uM) and NBQX (10 uM).

### Viral Vectors for Functional and Connectivity Studies

AAV8-hSyn-DIO-mCherry (Addgene, catalog # 50459) was used to determine the optimal injection coordinates for target Trh^Arc^, AAV1-hSyn-Flex-GCaMP6s (Addgene, catalog # 100845) was used for two-photon slice experiments to record calcium signaling in Trh^Arc^ neurons following application of liraglutide or leptin, pAAV-EF1alpha-dFlox-hChR2-mCherry (Addgene, catalog # 20297) was used for CRACM experiments evaluating the monosynaptic connection from Trh^Arc^ to AgRP arcuate neurons, pAAV-EF1alpha-dFlox-hChR2-eYFP (Addgene, catalog # 20298) was used for all optogenetic experiments and anterograde experiments, AAV-hSyn-fIO-hM4Di-mCherry and AAV-DJ-CMV-DIO-EGFP-2A-TeNT (gift from Richard Palmiter) was used for experiments testing the necessity of Trh-Cre+ neurons for regulating body weight, food intake, and responses to liraglutide, leptin, and saline injections.

### Rabies Injections for RAMPANT experiments

Agrp-Cre heterozygous mice were anesthetized with ketamine (80 mg/kg) and xylazine (10 mg/kg) diluted in phosphate-buffered saline (PBS). Mouse body temperature was maintained at 37 °C throughout the surgery with a closed loop infrared warming system (RightTemp Jr. or SomnoSuite, Kent Scientific). Once anesthetized, the mouse’s head was secured between the ear bars of a stereotaxic apparatus (Kopf). Local analgesic was provided by long-acting Bupivacaine (50-100 nL; Nocita). After waiting 5 minutes for local analgesia effect, skin overlying the skull was incised and retracted to expose the skull surface. Next, a small craniotomy was drilled above the Arc. A glass micropipette and Nanoject III microinjection system (Drummond Scientific) was used to inject into the bilateral Arc an adeno-associated virus (AAV) vector (AAV8-hSyn-FLEX-TVA-P2A-eGFP-2A-oG; titer, 3.10 x 10^13^ transduction units (TU)/mL; Salk Institute for Biological Studies; 400 nL total; 100 nL per injection; rate: 60 nL/second). Arc injection coordinates were based on “The Mouse Brain in Stereotaxic Coordinates” ^93^: AP: –1.48 mm, –1.62 mm, DV: –5.8 mm, ML +/− 0.2 mm. The pipette was slowly withdrawn from the injection site 5 minutes after each injection to avoid backflow. The incised scalp was sutured closed with surgical glue (Vetbond). Mice were provided with Meloxicam Sustained-Release (ZooPharm; 5mg/kg; IP) for post-operative analgesia, 1 mL of lactated Ringers solution in 5% dextrose to support hydration, and returned to the vivarium once ambulatory. Three weeks after AAV injections, mice again underwent stereotactic surgery to receive bilateral injections of EnvA-rabies-deltaG-H2b-mCherry into the Arc (400 nL total; coordinates above; titer: 7.35 x 10^9^ TU/mL; Salk Institute for Biological Studies). After recovery, mice were returned to their home cage and allowed to recover for 5 days before brain tissue was harvested.

### AAV-DIO-H2b-mCherry Injections

A separate cohort of Agrp-Cre mice (n=5; 4 male and 1 female; mean age: 13 weeks old +/− 8 weeks) underwent stereotactic surgery as described above, except to inject AAV9-DIO-H2b-mCherry (plasmid gifted by Dr. Bradford B. Lowell) into the bilateral ARC (400 nL total; coordinates above; titer: 3.94 x 10^13^ genome copies (GC)/mL; Vigene). After recovery, mice were returned to their home cage and allowed to recover for 3 weeks before tissue was harvested for sequencing.

### Feeding Conditions for RAMPANT experiments

Five days after rabies injection, each mouse underwent one of three feeding conditions: *ad libitum* feeding (“fed”), restricted from food for 13 hours (“fasted”), or restricted from food for 12 hours and then given *ab libitum* access to food for 2 hours (“post-fast re-fed”). Tissue harvest was performed at approximately the same time of day (+/− 1 hour) to minimize circadian effects on gene expression. All mice were allowed *ad libitum* access to water for the entirety of the experiment. Body weight of each mouse was documented before and after fasting and re-feeding to assess weight loss in fasted mice and weight regain in post-fast re-fed mice (Fig. S6A-C).

### Single-Nuclei RNA-Sequencing

Mice were rapidly decapitated without anesthesia for brain extraction to avoid stress and anesthesia-related changes in nuclear mRNA. Brains were immediately extracted and coronally sectioned at 500 μm intervals through the hypothalamus (Bregma +0.14 mm to –2.92 mm) using a Compresstome (VF-200-0Z Legacy Compresstome; Precisionary Instruments). Brain sections were immediately immersed in ice-cold RNAprotect reagent (Qiagen, catalog # 76106) to preserve RNA. Brain sections remained in RNAprotect in the dark overnight at 4 °C. The following day, brain sections were visualized under a fluorescence stereomicroscope (Zeiss Discovery V8) and regions of interest visibly containing H2b-mCherry+ nuclei were microdissected into chilled microcentrifuge tubes. Tissue samples were then stored at –80 °C for no longer than 2 weeks.

Once all samples for each batch were collected, tissue samples were thawed on ice and pooled by region and feeding condition. Tissue was dounce homogenized and purified by density gradient centrifugation into a single-nuclei suspension as previously described ^26,94^, but with the following modifications: in batch 2, TruSeq anti-Nuclear Pore Complex Proteins Hashtags were applied to track the brain region of origin for each cell nuclei (BioLegend, catalog # 682205, 682207, 682209). We counterstained nuclei with a far-red DNA fluorescent intercalator, DRAQ5 (Thermo Fisher, catalog # 62251, 1:500 dilution in sample) and kept them chilled on ice until sorting. We sorted single, DRAQ5+/mCherry+ nuclei using the Becton Dickenson FACS Aria Fusion (sample batch 1) or Becton Dickenson Influx (sample batch 2) cell sorters. All sorts were performed with SCYM (ASCP) certified technical assistance at the University of Virginia Flow Cytometry Core, using an 85 μm nozzle and set to purity mode. To separate nuclei from non-nucleated debris, we gated for events with high relative intensity of DRAQ5 fluorescence, using a 640 nm excitation and a 670/30 nm collection filter. We then gated the nuclei by forward scatter area *vs*. side scatter area to eliminate large aggregates, followed by forward scatter area *vs*. forward scatter height to select for singlets. Finally, from the single nuclei, we selected mCherry+ nuclei based on their high relative fluorescence, excitation 561 nm and 610/20 nm collection filter.

We sorted mCherry+ nuclei into 18.8 μL of RT Reagent B from the 10x Genomics Chromium Next GEM Single Cell 3’ Kit v3.1, added the remainder of the kit Step 1 master mix reagents to the capture tube, plus enough resuspension buffer to reach a total volume of 75 μL. We then processed the sample into complementary DNA (cDNA) sequencing libraries according to the manufacturer’s instructions (10X Genomics, CG000204 Rev D). Hashtag oligonucleotide (HTO) libraries were prepared according to a previously published protocol ^95^, with the following modification: double-sided SPRI (solid phase reversible immobilization) was repeated for a total of two times to cleanly separate HTO libraries from cDNA libraries after amplification and, if needed, just prior to pooling libraries for sequencing. Libraries were sequenced at a concentration of 1.85 pM and with a 75-cycle, high-output kit on the Illumina NextSeq 550 according to the manufacturer’s instructions.

### Sequencing Data Processing and Analysis

The 10X Genomics Cell Ranger pipeline (version 5.0) was used to map reads to the mouse reference transcriptome (mm10-2020-A) and quantify Unique Molecular Identifier (UMI)-corrected, gene-level expression values. Introns were included using the Cell Ranger include-introns parameter. The Cellbender software package was used to mitigate the effects of contaminating ambient RNA on our analysis ^96,97^. For libraries containing HTOs, the Seurat HTODemux() function was used to match each single nuclei transcriptome to a brain region-specific HTO. snRNA-seq feature-barcode matrices were analyzed in R (version 4.2.3) with Seurat v4.3.0 package ^98^.

We applied filters to each library based on their library specific distribution of quality metrics (parameters shown in Fig. S2). Libraries containing HTOs were filtered to remove any suspected doublets and HTO-negative cells. All 4 libraries were then integrated to correct for technical variance including batch effects ^98^. In brief, we log-normalized the data, selected 2,000 most variable genes for each batch (“feature selection”); integrated the libraries using the IntegrateData() function in Seurat; scaled each gene; performed Principal Component Analysis (PCA) to reduce linearly the dimensionality of the highly variable gene space; clustered the cells using the Louvian algorithm, based on Euclidean distance in the PCA space comprising the first 50 principal components (PCs) with a resolution value of 1.0; and performed non-linear dimensionality reduction by Uniform Manifold Approximation and Projection ^99^ for visualizing the clustered data in two dimensions. Cluster relatedness in PCA space was illustrated with dendrograms using the BuildClusterTree() function in Seurat.

Cell clusters were separated into three brain region-specific datasets according to each cluster’s HTO content. Clusters containing 5% or more cells labeled with a regional HTO were included in the corresponding regional dataset, accordingly. Clusters with more than one regional HTO, potentially representing cells near regional borders, were expected as an artifact of dissection. Each regional dataset was then re-clustered, including the steps of feature selection, PCA, and UMAP visualization. PC and resolution settings for each regional dataset are described in Supplemental Figure 2. Supervised cell-type annotation ^100^ was performed for Arc cell clusters using cell type labels from a previously published databases of hypothalamic neuron molecular subtypes ^30^. For this, a weighted vote classifier derived from the reference cell identities was used to predict cell identities for each rabies+ cell using Seurat’s FindTransferAnchors() function. Following integration, Seurat’s TransferData() function was used to transfer cell type labels from the Arc-ME reference dataset ^28^ and calculate cell-type prediction scores for each Arc rabies+ cell. Prediction scores are values between 0 to 1 which reflect the confidence of each cell-type prediction. Cells with prediction scores <0.5 were excluded from further analysis.

HypoMap labels were assigned by mapping rabies+ cells to the full hypoMap dataset using the mapscvi guided tutorial available online (https://github.com/lsteuernagel/mapscvi). Labels established at the C286 granularity were used for this analysis. Cells with prediction scores <0.5 and clusters with <10 cells were excluded from further analysis.

### RNA Fluorescence *In Situ* Hybridization (RNA FISH)

RNA FISH experiments were performed on brain tissue from mice that underwent monosynaptic rabies tracing from Agrp-Cre cells or from C57Bl6j mice following one of the three feeding conditions (see *Feeding Conditions for RAMPANT experiments*). Mice were terminally anesthetized with ketamine (80 mg/kg) and xylazine (10 mg/kg) diluted in PBS, followed by transcardial perfusion with 0.9% saline plus heparin and 4% paraformaldehyde (Thomas Scientific). Brains were extracted and post-fixed for 24 hr at 4 °C. Following fixation, brains were sectioned coronally at 35 μm thickness on a vibratome (Leica VT1000S).

The day before RNA FISH, sections were rinsed in PBS and then mounted on slides (Fisher Scientific) and left to dry overnight. An ImmEdge Hydrophobic Barrier Pen (Vector Laboratories) was used to draw a barrier around the sections. The sections were then incubated in Protease IV in a HybEZ II Oven for 30 min at 40 °C, followed by incubation with the target probe (*Agrp, Glp1r, Fos, Slc6a3*, *Ghrh*, *Trh*, or *Pomc*) for 2 hr at 40 °C. Slides were then treated with AMP 1– 3, HRP-C1, HRP-C2, HRP-C3, and HRP Blocker for 15–30 min at 40 °C, as previously described ^22^. TSA Plus FITC and TSA Plus Cy5 (Akoya Biosciences) were used for probe visualization. Rabies-H2b-mCherry was visualized using immunofluorescence with a rabbit anti-RFP primary antibody (Rockland, catalog # 600-401-379, 1:1000) overnight at room temperature and a donkey anti-rabbit 550 secondary antibody (Thermo Scientific, catalog # PISA510039, 1:1000) for 2 hr at room temperature.

Sections were coverslipped with mounting medium plus DAPI (Vector Laboratories, catalog # H-1200), and sealed with fingernail polish. Figure images were taken as Z-stacks using a confocal microscope (Zeiss LMS800). Optimal Z depth and slice interval were determined empirically for each image (20x, range of 6-14.85 μm, 4-12 slices; 63x, range of 3.6-8 μm, 5-31 slices). Z-stack images were compressed into a single image using ImageJ, with the projection type set to Maximum Intensity. For quantification, sections were imaged using a Revolve R4 fluorescence microscope (ECHO, RVL-100G). Anatomical borders were approximated based on a stereotactic atlas of the mouse brain ^76^.

### Virus Injections

Stereotaxic coordinates for Trh^Arc^ neurons were –2.20 AP, + − 0.23 ML, and –5.7 DV. During surgeries, mice were anesthetized with isoflurane, placed in a stereotaxic frame (Stoelting’s Just for Mouse), and provided with analgesia (Meloxicam, 0.5 mg/kg). Following a small incision on top of the skull and skull leveling, a small hole was drilled for injection. A pulled-glass pipette (20–40 mm tip diameter) was inserted into the brain at coordinates aimed at the Arc (AP: −2.20, ML: + − 0.23, DV: –5.7), and 100 nL of virus was injected using a micromanipulator (Grass Technologies, Model S48 Stimulator, 25 nL/min). For in vivo optogenetic experiments targeting Trh^Arc^ terminals and AgRP soma in the arcuate, the virus was injected to target Trh^Arc^ neuron somas (AP: –2.20, ML: + − 0.23, DV: –5.7) and AgRP neuron somas (AP: –1.5, ML: + − 0.23, DV: –5.7). The pipette was pulled up 10 minutes after injection to reduce the backflow of the virus.

### Optical Fiber Implantation

For *in vivo* optogenetic experiments targeting the Trh^Arc^ soma, an optic-fiber cannula (200um diameter core; catalog # CFMLC22U-20, Thor Labs) was implanted directly over the Arc (AP: −2.20, ML: + 0.3, DV: −5.60) following virus injection and fixed to the skull (C&B-Metabond Quick Adhesive Cement dental acrylic). For *in vivo* optogenetic experiments targeting AgRP neuron somas and/or Trh^Arc^ terminals, an optic fiber was placed directly over AgRP neuron somas (AP: –1.5, ML: + − 0.23, DV: –5.6). After recovery, mice were singly housed and allowed to recover for >3 weeks before further experimentation.

### Optogenetic Behavior Paradigm

For all optogenetic experiments, mice were habituated to the photostimulation setup and the FED3 device 2-3 times. Fiber optic cables (200 mm diameter, Doric Lenses) coupled to lasers were attached to the fiber cannula of the mice via zirconia sleeves (Doric Lenses). Light was delivered to the brain through an optical fiber (200 μm diameter core; CFMLC22U-20, Thor Labs). Light power exiting the fiber tip was 10 mW for all optogenetic experiments.

For photostimulation, pulse trains (20 Hz; 2 sec on, 2 sec off; 473 nm from Laserglow laser technologies) were programmed using a waveform generator (PCGU100; Valleman Instruments) for continuous photostimulation during all tasks. For simultaneous photoactivation of Trh^Arc^ neurons and chemogenetic inhibition of Npy^Arc^/AgRP neurons, in addition to optoegenetically targeting Trh^Arc^ soma, we injected a flp-dependent hM4Di DREADD targeting the rostral Arc (AP: –1.5, ML: + − 0.23, DV: –5.6) into Trh-Cre;Npy-FlpO mice. For fasted photostimulation experiments, mice were fasted for ∼18 hours overnight, and photostimulation experiments were conducted at 08:00–11:00 hours, near the beginning of the light cycle when food intake was low. For sated photostimulation experiments, photostimulation experiments were conducted at the onset of the dark cycle when food intake was high. For priming experiments, mice were stimulated for 1 hour before food presentation, stimulation was turned off, and food was presented.

### Food Intake from Feeding Experimentation Devices (FED3)

Feeding information was collected using FED3 devices ^54^, which dispense 20 mg pellets of chow food *ad libitum*. In all experiments using the FED3 device, mice were habituated to the device in the homecage for at least one day. Mice were considered trained on the FED3 when they consumed at least 200 pellets in a day (4 grams).

### Loss-of-Function Studies with Tetanus Toxin

Mice were single-housed and separated into two groups: Trh-Cre mice injected with Cre-dependent AAV-TeNT (tetanus toxin) and C57BL/6J mice injected with the same AAV, both with standard housing conditions and *ad libitum* access to food and water unless otherwise specified.

For experiments involving acute and chronic injections of liraglutide, or vehicle, both groups of mice were initially matched for age and weight, and habituated three times to the protocol before experiments began. For acute injections, the protocol consisted of removing the standard diet from each cage 3 hours before the onset of the dark cycle, injecting saline or liraglutide 30 minutes before the onset of the dark cycle, and reintroducing a FED3 device set to ad-libitum pellet mode at the onset of the dark cycle. Baseline measurements were taken after surgery but before viral expression had occurred, and experimental measurements were collected approximately 8 weeks after surgery. For the chronic injection experiments, wildtype mice were given ad-libitum access to HFD to bring their bodyweight to comparable levels as the Trh^Arc^-TeNT mice, which were also acclimated to HFD. Baseline body weight and food intake (chow and HFD) measurements were taken over five days before the start of the protocol. After this baseline period, weight and food intake were measured for one week during which concurrent liraglutide injections were administered.

The same mice injected with liraglutide and vehicle were also used to measure the long-term effects of TeNT on feeding and body weight. Baseline measurements were taken after the mice had been singly housed for 1 week. Six baseline measurements of body weight and food intake were averaged together per mouse. Experimental measurements began three weeks after AAV injection surgery.

### *Ex Vivo* Calcium Imaging

Trh-Cre mice of age 2-3 months were injected with AAV1-hSyn-Flex-GCaMP6s (Addgene, catalog # 100845). More than 3 weeks after surgery, mice were anesthetized by isoflurane and decapitated. The brain was quickly extracted and immediately placed into ice-cold, carbogen-saturated (95% O_2_, 5% CO_2_), choline cutting solution (in mM): 2.5 KCl; 1.25 NaH_2_PO_4_; 20 HEPES; 10 MgSO4.7H20; 0.5 CaCl_2_; 92 Choline Chloride; 25 Glucose; 2 Thiorea; 5 Sodium Ascorbate; 3 Sodium Pyruvate; and 30 NaHCO_3_ (pH 7.3 – 7.6). Then, 275 μm thick coronal sections of the Arc were cut with a Campden Instruments 7000smz-2 Vibratome. Slices were settled in the same carbogen-saturated cutting solution for 10 minutes at 36 °C, and then incubated at 36 °C in oxygenated aCSF for 60 minutes (in mM): 125 NaCl; 21.4 NaHCO_3_; 2.5 KCl; 1.2 NaH_2_PO_4_; 1.4 CaCl_2_; 10 Glucose; and 1.2 MgCl_2_. Slices were maintained and recorded at room temperature (20-24 °C) until transferred to an immersion-recording chamber and superfused at a rate of 2.5 mL min^-1^.

Two-photon imaging was performed using a multiphoton laser scanning microscope (Olympus FVMPE-RS) at 33.3 frames/second and 512 x 512 pixels/frame as described previously ^101^. An InSight X3 laser (Spectra-Physics) was used to excite the fluorophore (920 nm), and the emission light was filtered (green: 495 – 540 nm) before collection with a GaAsP photomultiplier tube. The XY scanning was performed using resonant/galvo mirrors, and the Z scanning was performed by motorized z-axis (average slice number, 27; step-size, 5 μm; average cycle rate, 31.73 sec). For two-photon imaging of acute brain slices, slices were transferred to a recording chamber perfused with aCSF (oxygenated with 95% O_2_ and 5% CO_2_; flow rate: 2–5 mL/min) at room temperature. Imaging was performed with a 20x 1.0 NA water-immersion objective (Olympus). The excitation wavelength used was 920 nm.

### Data Processing and Analysis for *Ex Vivo* Calcium Imaging

Following data acquisition, Olympus Fluoview files (.oif) were opened in Fiji (ImageJ). A grouped z-project was used to obtain the max intensity across z-stacks, rigid body registration was used to reduce motion artifacts (TurboReg), and then the z-project was used to get an average intensity projection over the course of the recording. This max z-projection was exported to Cellpose2.0 ^102^ (Mouseland), and the “cyto” model was used to automatically select ROIs. Based on the model performance, ROIs were manually added or deleted. ROIs were exported back to Fiji, and the change in fluorescence was calculated by subtracting the 5 minute baseline fluorescence (F0(t)) from F(t), then dividing by F0(t): DF/F(t) = (F(t) – F0(t))/F0(t) for each ROI. To calculate “responsive” cells, the Z-score for each ROI was calculated by subtracting the mean change in fluorescence during the 5-min baseline period from F(t) and dividing by the standard deviation of mean fluorescence from the 5-min baseline period. Cells were “responsive” if the average Z-score during the 10-min wash on period of either saline, liraglutide, or leptin was above or below 1.645. These results were exported to Prism for further analysis.

### *Ex Vivo* Whole-cell Patch Clamp Electrophysiology

Channelrhodopsin (ChR2)-assisted circuit mapping (CRACM) was performed. CRACM involves *in vivo* targeted expression of ChR2, a photo-excitable cation channel, in presumptive presynaptic upstream neurons (and their terminals), followed by *ex vivo* electrophysiologic assessment in acute brain slices of light-evoked postsynaptic currents in candidate downstream neurons. Trh-Cre;Npy-hrGFP received bilateral 100 nL injections of AAV9-EF1alpha-dFlox-hChR2-mCherry in the Arc. Four to 6 weeks later, brain slices were obtained and stored at 30 °C in a heated, oxygenated chamber containing aCSF (in mmol/L) 124 NaCl, 4 KCl, 2 CaCl2, 1.2 MgSO4, 1 NaH2PO4, 10 glucose, and 26 sodium bicarbonate before being transferred to a submerged recording chamber maintained at 30 °C (Warner Instruments, Hamden, CT). Recording electrodes (3-5 MOhm) were pulled with a Flaming-Brown Micropipette Puller (Sutter Instruments, Novato, CA) using thin-walled borosilicate glass capillaries. Light-evoked inhibitory postsynaptic currents (IPSCs) were measured in voltage-clamp mode using electrodes filled with an intracellular recording solution containing (in mM): 130 CsCl, 1 EGTA, 10 HEPES, 2 Mg-ATP, 0.2 Na-GTP. To isolate GABAergic synaptic transmission, kynurenic acid (3 mM) was included in the bath aCSF, and the Npy GFP+ neurons held at –70 mV. Light evoked IPSCs was recorded in the presence of Tetrodotoxin (TTX, 500 nM) and 4-aminopyridine (4-AP, 100 mM).

### Rabies Injections for Trh^Arc^ Neuron Tracing

Monosynaptic rabies tracing was performed in Trh-Cre mice (n=3, age 2-3 months) using the same viruses described above. Brain tissue was harvested 7 days after injection with [EnvA]rabies-H2b-mCherry. Anatomical borders for quantified regions were established using a stereotactic atlas of the mouse brain ^93^.

### Perfusion and Histology

After completing behavioral experiments, mice with viral injections and/or optical implants were terminally anesthetized using chloral hydrate (Sigma-Aldrich, catalog # 301-17-0) and transcardially perfused first with phosphate-buffered saline (PBS) followed by 10% neutral buffered formalin (Fisher Scientific, catalog # SF100). Brains were removed, post-fixed, and dehydrated in 30% sucrose before sectioning into 30–50 um slices using a freezing sliding microtome (Leica Biosystems). Coronal sections were collected and stored at 4 °C. Slices were mounted with a mounting medium containing DAPI (Vectashield, catalog # H-1200-10), and images were captured using a 10X objective on an Olympus VS200 Scanscope and 20x objective on a Zeiss Observer Z1 confocal microscope.

### Statistical Analysis

GraphPad Prism 10 was used for statistical analysis, and GraphPad Prism10 and Adobe Illustrator 2020 were used to generate graphs. For discrete comparisons between two groups, two-tailed Student’s t-tests were used. For comparisons across groups or between groups over time, repeated measures one-way or two-way ANOVAs were used, respectively, with corresponding post hoc tests adjusted for multiple comparisons. Normality and equal variances were assumed. Mice were randomly assigned to groups but were matched for age, sex, and body weight. Except for the RNA FISH quantitative analysis, experimenters were not blinded to conditions during testing and analysis. Power analyses were not used to determine sample sizes, however, group sizes were chosen to match similar studies.

Key resources table.

**Key resources table**
| REAGENT<br>or<br>RESOURCE | SOURCE | IDENTIFIER |
| --- | --- | --- |
| Antibodies |  |  |
| anti-RFP<br>primary<br>antibody | Rockland | 600-401-379 |
| anti-rabbit<br>550<br>secondary<br>antibody | Thermo<br>Scientific | PISA510039 |
| TotalSeq™<br>-A0451<br>anti-Nuclear<br>Pore<br>Complex<br>Proteins<br>Hashtag 1<br>Antibody | BioLegend | 682205 |
| TotalSeq™<br>-A0452<br>anti-Nuclear<br>Pore<br>Complex<br>Proteins<br>Hashtag 2<br>Antibody | BioLegend | 682207 |
| TotalSeq™<br>-A0453<br>anti-Nuclear<br>Pore<br>Complex<br>Proteins<br>Hashtag 3<br>Antibody | BioLegend | 682209 |
| Bacterial and virus strains |  |  |
| AAV8-hSyn-FLEX-TVA-P2A-eGFP-2A-oG | Salk Institute<br>for Biological<br>Studies<br>custom<br>preparation | Not applicable |
| EnvA-rabies-deltaG-H2b-mCherry | Salk Institute<br>for Biological<br>Studies<br>custom<br>preparation | Not applicable |
| AAVDJ-DIO-H2b-mCherry | Vigene custom<br>preparation | Not applicable |
| AAV8-hSyn-DIO-mCherry | Addgene | 50459 |
| AAV1-hSyn-Flex-GCaMP6s | Addgene | 100845 |
| pAAV-EF1alpha-dFlox-hChR2-mCherry | Addgene | 20297 |
| pAAV-EF1alpha-dFlox-hChR2-eYFP | Addgene | 20298 |
| AAV-hSyn-fIO-hM4Di-mCherry | gifted from<br>Richard<br>Palmiter | Not applicable |
| AAV-DJ-CMV-DIO-EGFP-2A-TeNT | gifted from<br>Richard<br>Palmiter | Not applicable |
| Chemicals, peptides, and recombinant proteins |  |  |
| Leptin | R & D Systems | 498-OB |
| Liraglutide | Selleckchem | S8256 |
| KCl | Sigma Aldrich | 7447-40-7 |
| Picrotoxin | Abcam | ab120315 |
| NBQX | Abcam | ab120046 |
| DL-AP5 | Abcam | ab120004 |
| TTX | Abcam | Ab120055 |
| 4-AP | Abcam | Ab120122 |
| TSA Plus<br>Cy5 | Akoya<br>Biosciences | NEL745001KT |
| TSA Plus<br>FITC | Akoya<br>Biosciences | NEL741001KT |
| Critical commercial assays |  |  |
| RNAscope<br>Multiplex<br>Fluorescent<br>Reagent Kit<br>v2 | Advanced Cell<br>Diagnostics | 323110 |
| RNAscope<br>H2O2 and<br>Protease<br>Reagents | Advanced Cell<br>Diagnostics | 322381 |
| Chromium<br>Next GEM<br>Chip M<br>Single Cell<br>Kit | 10X Genomics | 1000371 |
| Deposited data |  |  |
| Raw and<br>processed<br>single-nucli<br>RNA-seq<br>data files | GEO | To be determined |
| Clustered<br>single<br>nuclei<br>RNA-seq<br>data | Broad Single<br>Cell Portal | To be determined |
| Experimental models: Organisms/strains |  |  |
| Mouse/Agr<br>p-IRES-Cre | Jackson<br>Laboratories,<br>ref. <sup>89</sup> | 012899;<br>RRID:IMSR_JAX:012899 |
| Mouse/Trh-IRES-Cre | Jackson<br>Laboratories,<br>ref. <sup>23</sup> | 032468;<br>RRID:IMSR_JAX:032468 |
| Mouse/Npy<br>-IRES2-FlpO-D | Jackson<br>Laboratories,<br>ref. <sup>90</sup> | 030211<br>RRID:IMSR_JAX:030211 |
| Mouse/Npy<br>-hrGFP | Jackson<br>Laboratories,<br>ref. <sup>48</sup> | 006417;<br>RRID:IMSR_JAX:006417 |
| Oligonucleotides |  |  |
| Mm-Slc6a3-C3 | Advanced Cell Diagnostics | 315441-C3 |
| Mm-Ghrh-C3 | Advanced Cell Diagnostics | 470991-C3 |
| Mm-Trh | Advanced Cell Diagnostics | 436811 |
| Mm-Pomc-C2 | Advanced Cell Diagnostics | 314081-C2 |
| Mm-AgRP | Advanced Cell Diagnostics | 400711 |
| Mm-Glp1r-C2 | Advanced Cell Diagnostics | 418851-C2 |
| Mm-Fos-C2 | Advanced Cell Diagnostics | 316921-C2 |
| Mm-Fos-C3 | Advanced Cell Diagnostics | 316921-C3 |
| Recombinant DNA |  |  |
| Software and algorithms |  |  |
| Cell Ranger v5.0 | 10x Genomics | https://support.10xgenomics.com/single-cell-gene-expression/software/pipelines/latest/what-is-cell-ranger; RRID:SCR_017344 |
| bcl2fastq v2.20.0 | Illumina | https://support.illumina.com/sequencing/sequencing_software/bcl2fastq-conversion-software.html RRID: SCR_015058 |
| R | version 4.2.3 | https://www.r-project.org/ RRID:SCR_001905 |
| Seurat v4.3.0 | Ref. <sup>100</sup> | https://github.com/satijalab/seurat RRID:SCR_016341 |
| Seurat code used from cell clustering | Zenodo | To be determined |
| Illustrator | Adobe | https://www.adobe.com; RRID:SCR_010279 |
| Excel | Microsoft | https://www.microsoft.com/en-us/; RRID:SCR_016137 |
| ImageJ | version 2.9.0 | https://imagej.net/software/fiji/ RRID:SCR_003070 |
| Mouseland Cellpose | version 2.2.2 | https://github.com/MouseLand/cellpose |
| FED3 Software | version 1.16.3 | https://github.com/KravitzLabDevices/FED3_library |
| Other |  |  |
| FED3 devices | Ref. <sup>64</sup> | https://github.com/KravitzLabDevices/FED3 |

## DATA AND CODE AVAILABILITY

The raw and processed snRNA-seq data are available at the Gene Expression Omnibus (GEO) repository at GEO accession number #####. The code for further processing and analyzing the data using Seurat can be found at Zenodo record number #####. The whole RAMPANT dataset (all-rabies) and its Arc-only subset are each available for user-friendly exploration through the Broad Single Cell Portal at these links: all-rabies, #####; Arc-only, #####. R data files (.RDS) of these fully processed and analyzed datasets can be downloaded directly from their Broad Single Cell Portal site. All other raw and processed data are available from the corresponding authors upon request.

## ACKNOWLEDGMENTS

We gratefully acknowledge technical contributions from Nicholas Conley and Ruei-Jen Abraham-Fan. We thank Jinhua Cang and Xiaorong Liu for the use of their confocal microscope and Cheng Zhan, Fuqiang Xu, and Minmin Luo for sharing data from their published study on rabies mapping of synaptic inputs to AgRP neurons ^22^. The rabies studies were supported in part by the GT3 Core Facility of the Salk Institute with funding from NIH-NCI CCSG: P30 CA01495, an NINDS R24 Core Grant and funding from NEI. Cell sorting and cytometry was performed by the University of Virginia Flow Cytometry Core Facility (RRID:SCR_017829), which is partially supported by a National Cancer Center award (P30 CA044579). Sequencing on the Illumina Next-Seq platform was performed by the Genomics Core of the Biology Department at University of Virginia and by the Genome Analysis and Technology Core of University of Virginia’s School of Medicine (RRID:SCR_018883). The study overall was funded by: a University of Virginia Brain Institute Fellowship to A.N.W.; American Diabetes Association Pathway to Stop Diabetes (Initiator Award 1-18-INI-14), National Heart, Lung, and Blood Institute (R01 HL153916), and National Eye Institute (R21 EY033528) awards to J.N.C.; Intramural Research Program of the National Institutes of Health and National Institute of Diabetes and Digestive and Kidney Diseases (DK075088 and DK075087-06) awards to M.J.K.. T.H.P. acknowledges the Novo Nordisk Foundation (unconditional donation to the Novo Nordisk Foundation Center for Basic Metabolic Research; grant number NNF18CC0034900) and the Danish Council for Independent Research (grant number 8045-00091B).

## CONTRIBUTIONS

A.N.W.; J.J.B.; N.H.; M.J.K.; and J.N.C. conceived and designed the study. A.N.W. performed the RAMPANT and RNA FISH studies, except for those of *Fos*/*Trh*/*Glp1r* and *Trh*/*Agrp*. J.J.B. performed the feeding/body weight studies, two-photon slice calcium imaging, rabies mapping of Trh^Arc^ neurons, and RNA FISH of *Trh* and *Agrp*. D.C.S. re-analyzed a previously published reference dataset of Arc neuron subtypes and with M.J. assisted in processing the RAMPANT data. D.C., E.O.K., E.G.D, and A.D. helped perform the functional studies. D.K. and A.L. trained J.B. on the two-photon slice calcium imaging protocol and assisted with experimental design and analysis. C.L. performed the electrophysiological connectivity studies. C.B., R.A.O., M.C., and M.J. performed RNA FISH of *Fos*/*Trh*/*Glp1r*. E.N.G. and A.D.G. contributed key methods for preparing tissue samples for RAMPANT. D.M.B.-R. and T.H.P contributed key methods for hashtagging nuclei and single-nuclei library preparation. A.N.W.; J.J.B.; C.L.; D.C.S; and C.B. performed the data analysis. A.N.W.; J.J.B.; M.J.K.; and J.N.C. wrote the paper with input from all authors.

## FIGURES

**Supplemental Figure 1.**
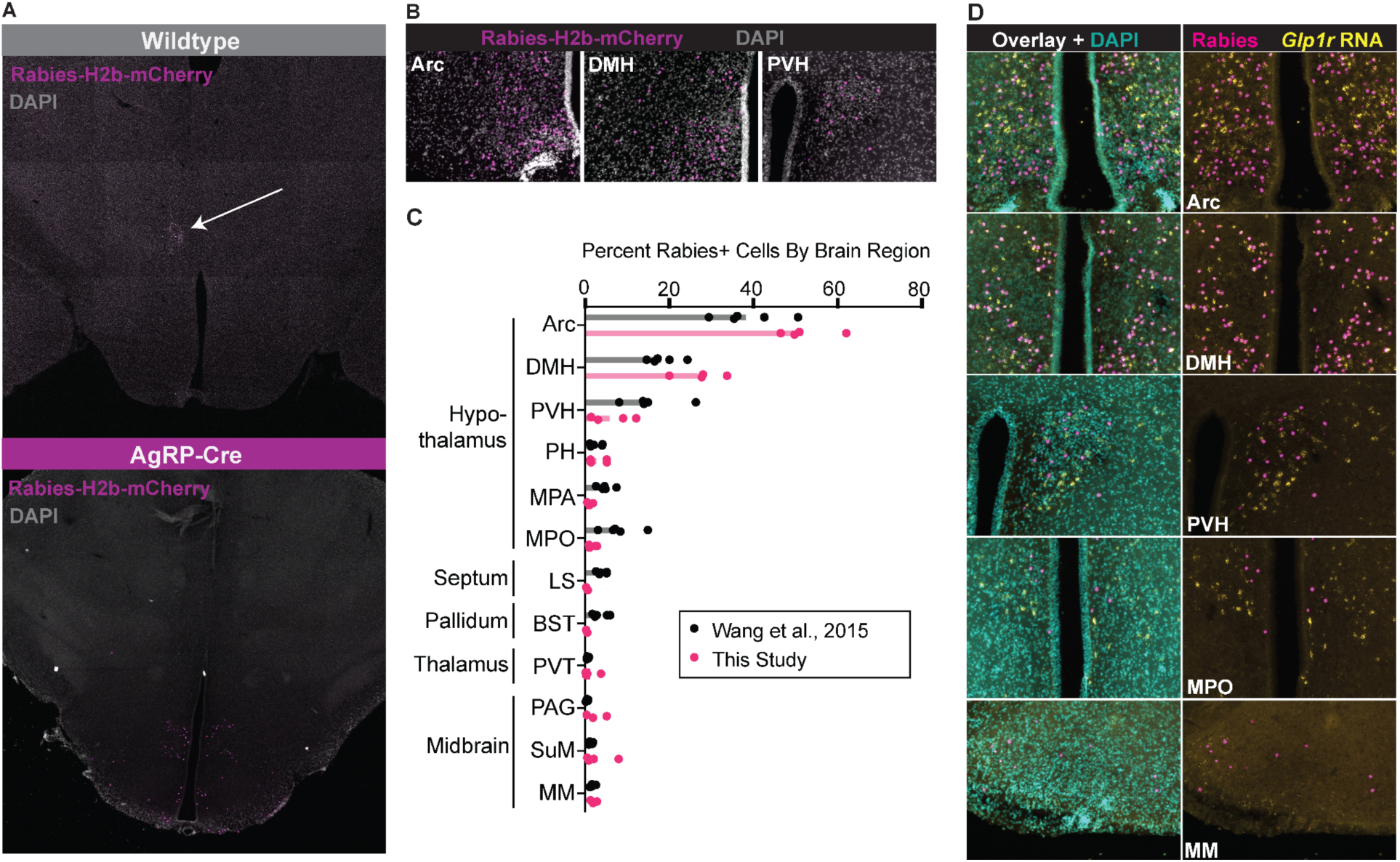
– Validation of Virus Specificity, Comparison with Previous Rabies Mapping Results, and Regional Identification of Glp1r+ Afferents to AgRP Neurons. **A**, Representative images from brains of C57BL6j mice (n=5) and Agrp-Cre mice after injection of Cre-dependent rabies helper AAV (AAV8-hSyn-FLEX-TVA-P2A-eGFP-2A-oG) followed by EnvA-rabies-deltaG-H2b-mCherry (“rabies-H2b-mCherry”). In C57Bl6j mice, both injections were targeted to the same site in the thalamus to ensure successful co-injection. Arrow indicates injection site. In Agrp-Cre mice, both injections were targeted to the Arc. **B,** Representative images of monosynaptic rabies labeling of AgRP neurons and their afferents in the arcuate hypothalamus (Arc), dorsomedial hypothalamus (DMH), and paraventricular hypothalamus (PVH) with rabies-H2b-mCherry. **C,** Comparing the present study and ref. ^22^ in terms of the percentage of cells labeled with rabies-H2b-mCherry via AgRP neurons in various brain regions (n=4 mice for this study, 5 mice for ref. ^22^). Bars represent means. MPA, medial preoptic area; MPO, medial preoptic nucleus; LS, lateral septum; BST, bed nucleus of the stria terminalis; PVT, paraventricular thalamus; PAG, periaqueductal gray; SuM, supramammillary nucleus; MM, medial mammillary nucleus. **D,** Representative images of *Glp1r* RNA FISH and monosynaptic rabies-H2b-mCherry labeling via AgRP neurons in several brain regions known to contain afferents to AgRP neurons and *Glp1r*+ neurons. The Arc and DMH contained the highest densities of rabies-labeled *Glp1r*+ afferents to AgRP neurons (n=4 mice).

**Supplemental Figure 2.**
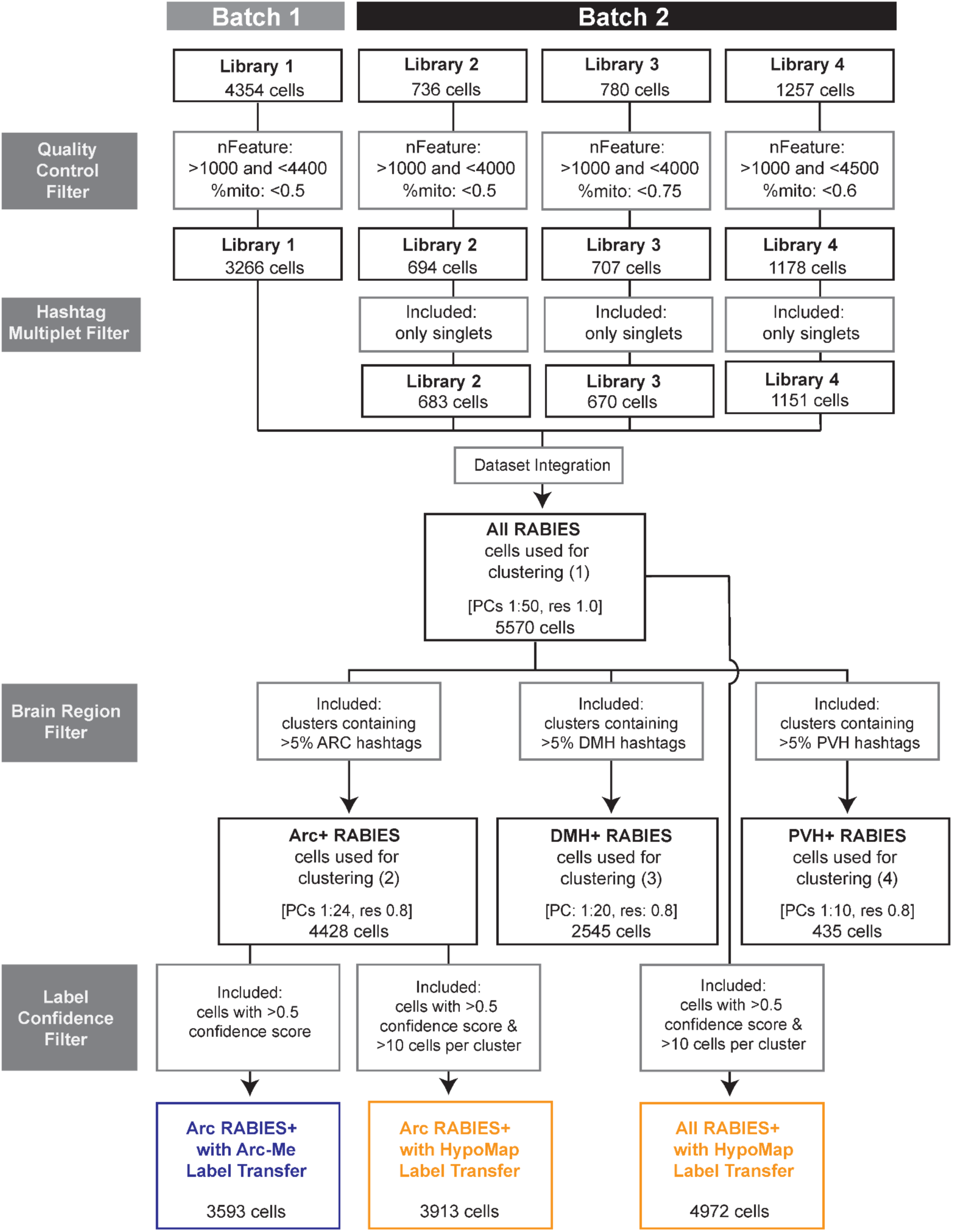
– Schematic of Data Processing Pipeline. Summary of filtering and Seurat cell clustering parameters. %mito, percent of reads from mitochondrial genes. PCs, principal components. Res, resolution. nFeature, number of unique genes detected. Hashtag, tissue sample-specific molecular barcode. Confidence, cell type prediction score based on label transfer from reference dataset.

**Supplemental Figure 3.**
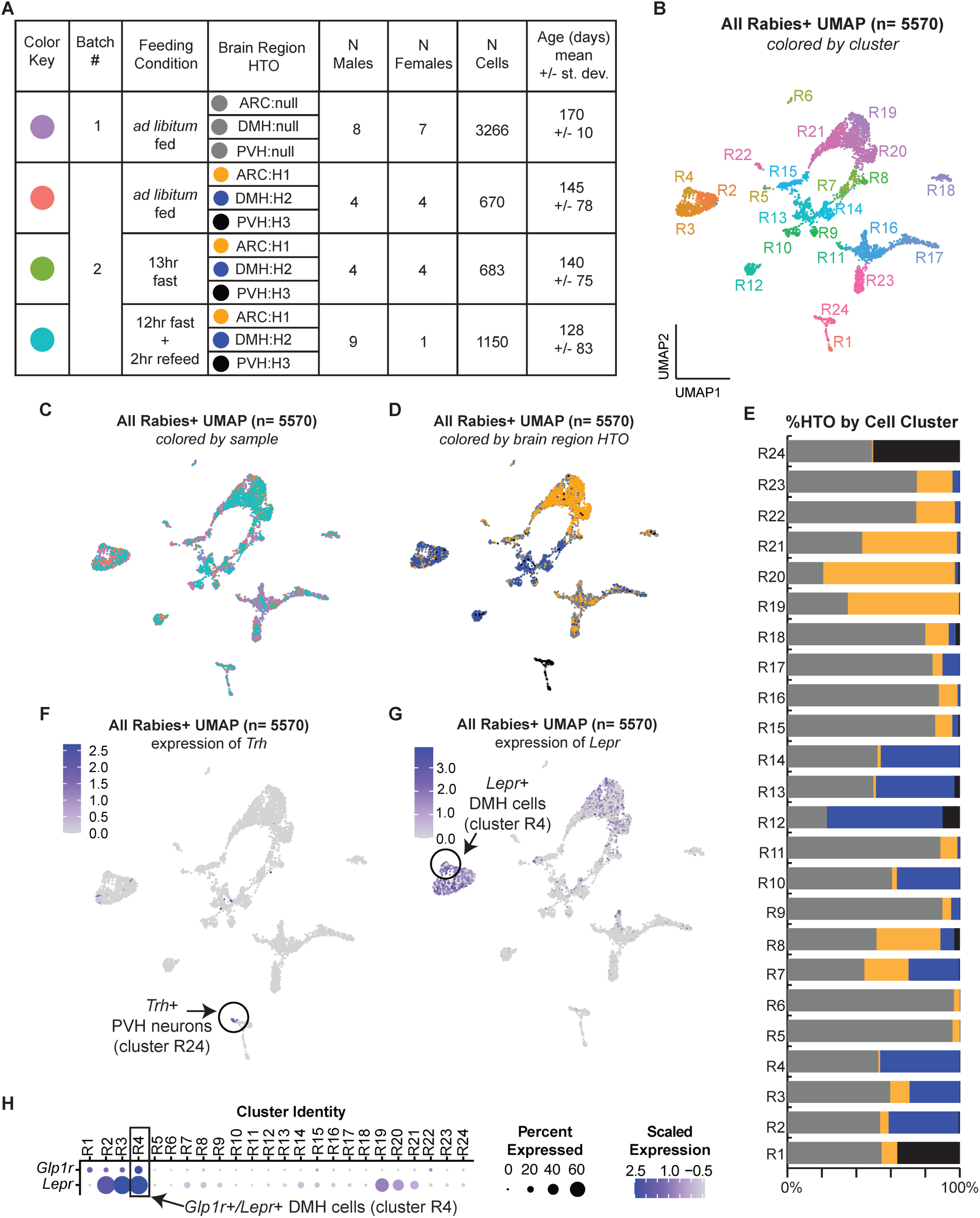
– Quality Control and Identification of Arc, DMH, and PVH Cells. **A**, Table of sample metadata and corresponding color labels (note, hashtag oligonucleotides, HTOs, were not used for batch 1). **B**, UMAP of all RAMPANT cells after initial quality control filtering, colored by cluster ID. **C**, Same UMAP as in panel A but colored according to sample (see panel A for color key). **D**, Same UMAP as in panel A but colored according to Arc, DMH, or PVH HTO. **E**, HTO composition of each cell cluster shown in panel a; batch 1 cells, which were not hashtagged and so lack HTOs, are indicated in gray. **F**, Same UMAP as in panel A but colored to indicate *Trh* gene expression. **G**, Same UMAP as in panel A but colored to indicate *Trh* gene expression. **H**, *Glp1r* and *Lepr* expression in all-rabies clusters.

**Supplemental Figure 4.**
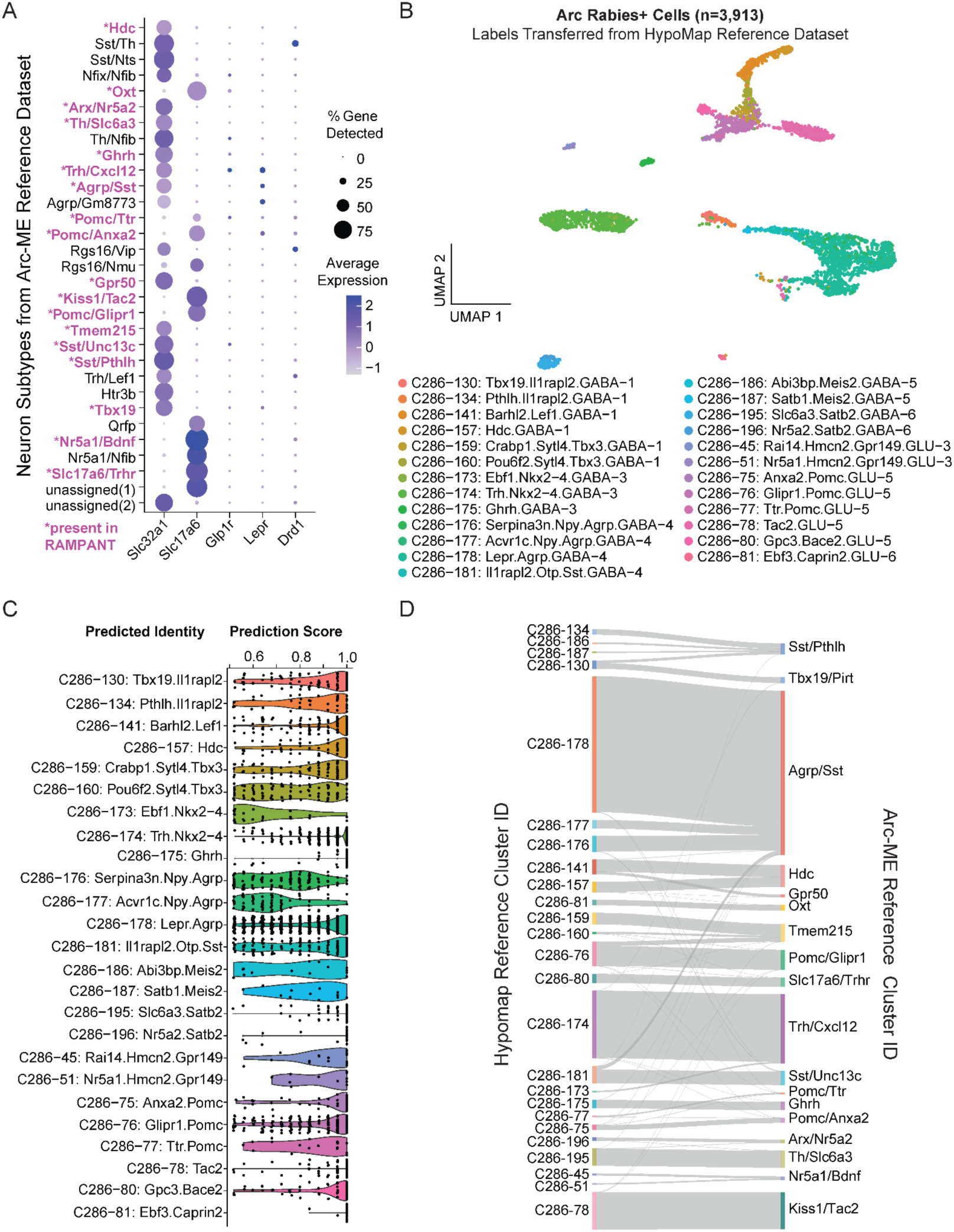
– Characterization and Identification of Arc Neuron Subtypes. **A**, Expression of select genes of interest among Arc neuron subtypes from a previous publication^30^. Neuron subtypes detected in RAMPANT analysis of AgRP neurons and their afferents are indicated in bold magenta. Dot size indicates the percentage of cells in that cluster in which the gene was detected, whereas the color represents the gene expression level after log normalization and scaling. **B,** UMAP of Arc rabies+ cells after transferring cell-type labels from HypoMap reference atlas of Arc neuron subtypes ^37^. **C,** Prediction score of transferred HypoMap cell-type labels after filtering out cells with low prediction scores (<0.5). **D,** Correspondence between cell-type labels transferred from HypoMap reference atlas and Arc-ME reference atlas.

**Supplemental Figure 5.**
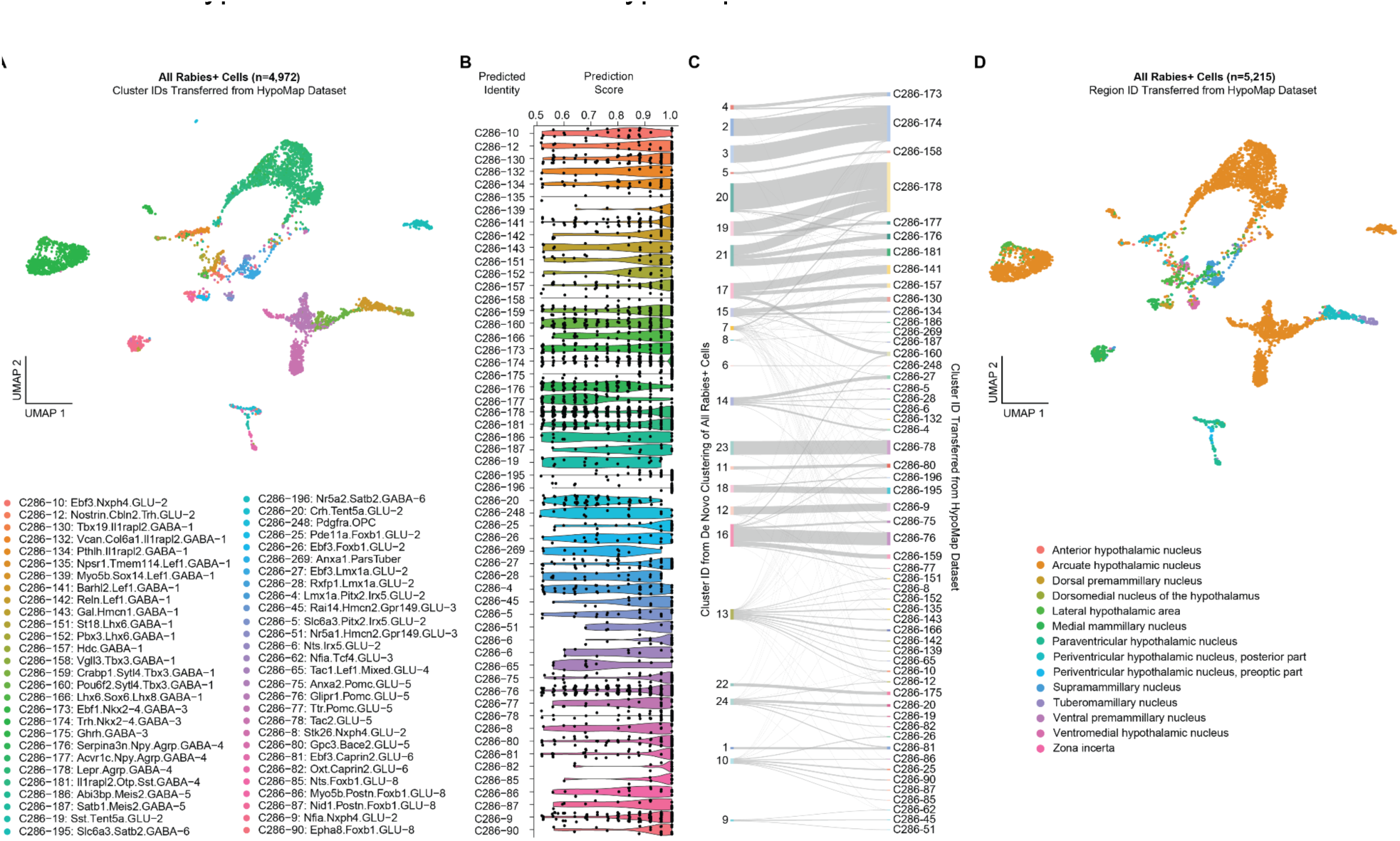
– Identification of Rabies-Infected Cells by HypoMap Label Transfer. **A**, UMAP of all rabies+ cells after transferring cell-type labels from the mouse HypoMap reference atlas ^37^ and filtering out cells with low prediction scores (<0.5). **B,** Prediction scores for assigning a cell type to each rabies+ cell by HypoMap label transfer. **C,** Correspondence between de novo clusters of all rabies+ cells and cell-type assignments from HypoMap label transfer. Each line represents a single cell, with its de novo cluster ID and HypoMap cluster ID on the left and right side, respectively. **D,** UMAP of all rabies+ cells after transferring regional labels from the mouse HypoMap reference dataset and filtering out cells with low prediction scores (<0.5).

**Supplemental Figure 6.**
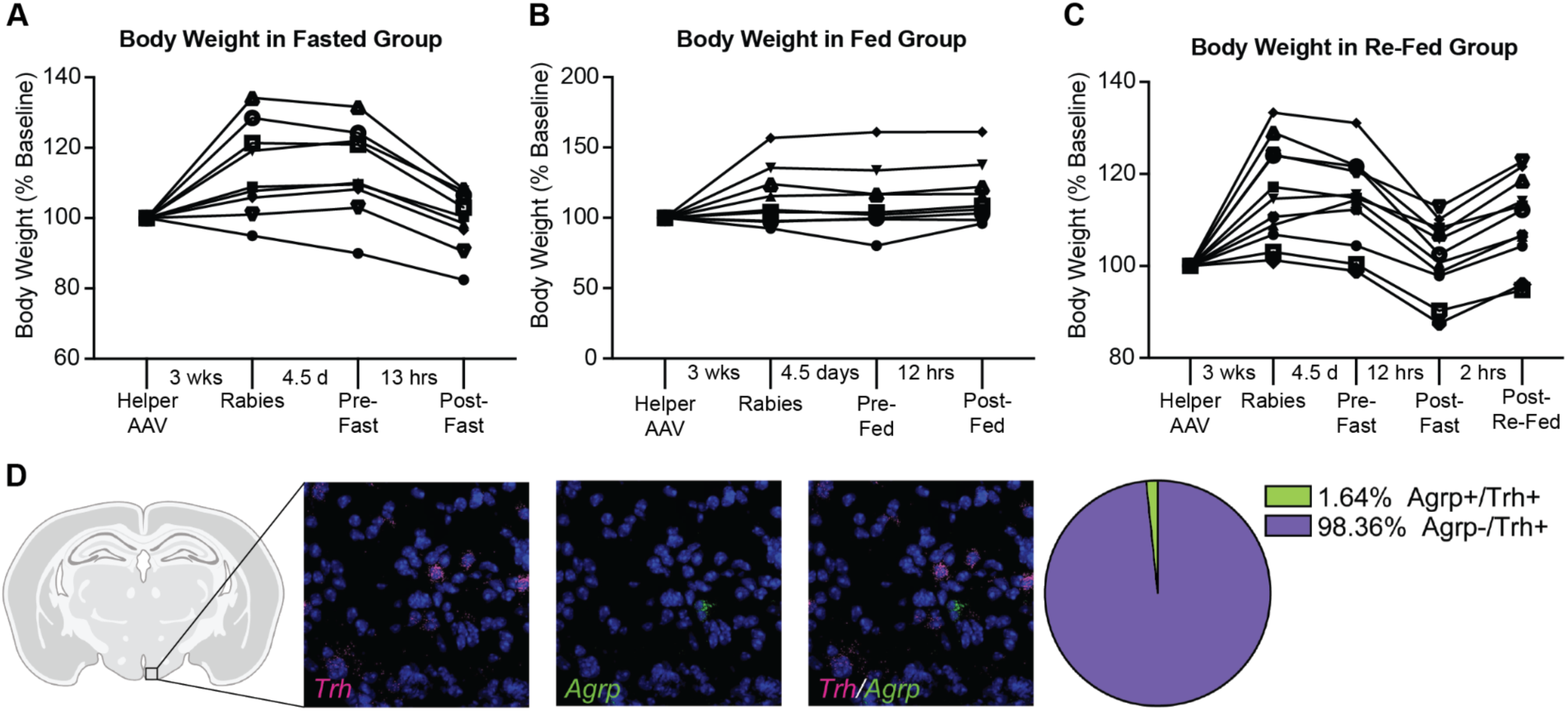
– Body Weights After Viral Injections and Feeding Conditions; Co-Localization of Agrp and Trh RNA in the Arc. **A**, Body weight in fasted group after injection of helper AAV and rabies, and before and after fasting. **B,** Body weight in *ad libitum* fed group after injection of helper AAV and rabies, and before and after night of *ad libitum* feeding. **C,** Body weight in post-fast re-fed group after injection of helper AAV and rabies, and before and after fasting and re-feeding. **D,** *Agrp* and *Trh* RNA FISH in the Arc indicates that AgRP neurons and Trh^Arc^ neurons are essentially distinct populations (n=289 cells from 4 mice).

**Supplemental Figure 7.**
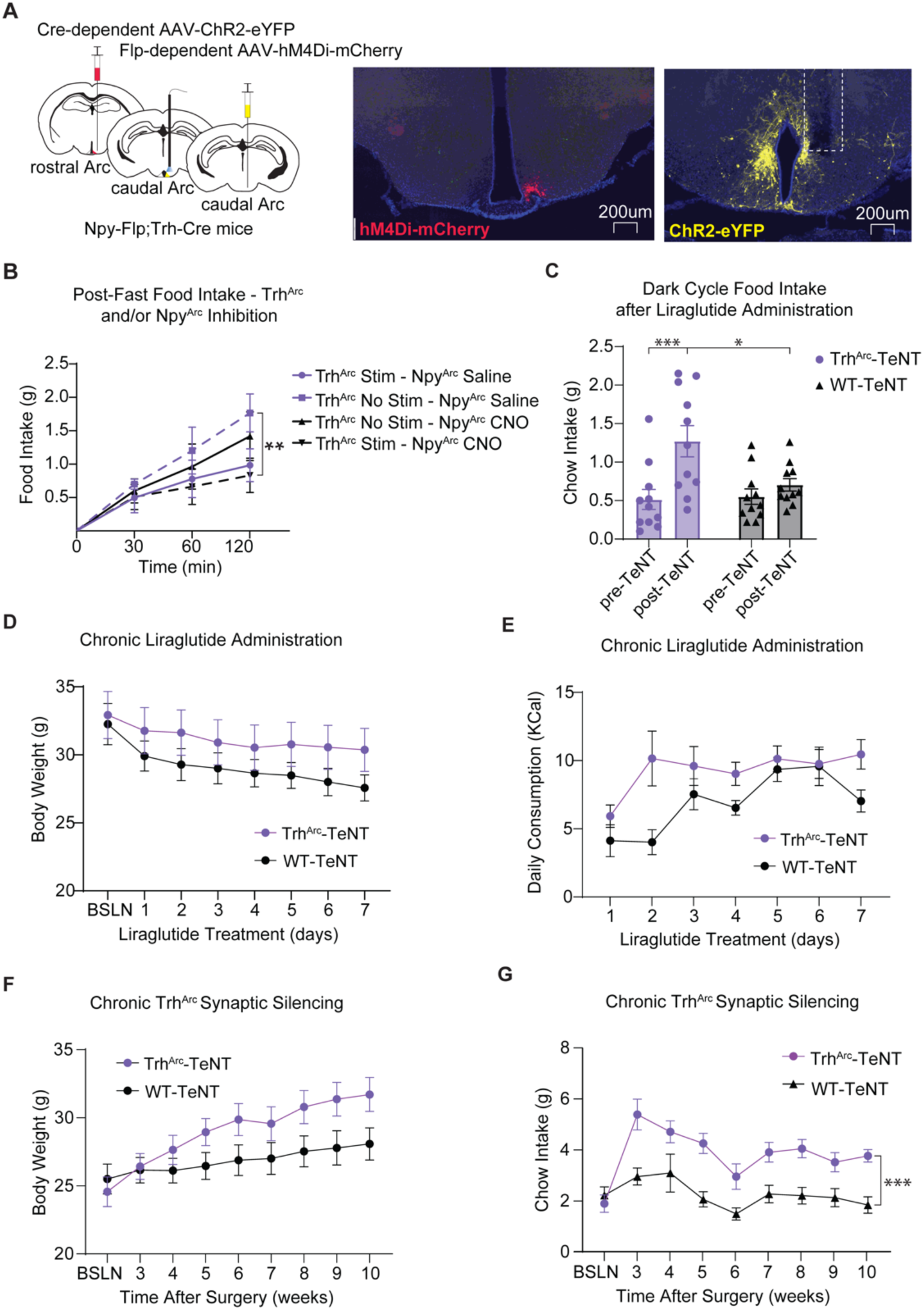
– Additional Occlusion and Loss-of-Function Studies. **A**, Left, schematic of unilateral injection of Cre-dependent AAV-ChR2 to caudal Trh^Arc^ neurons and optical fiber implant over the caudal Arc in Trh-Cre mice and unilateral injection of Flp-inducible AAV-hM4Di to rostral Npy^Arc^ neurons. Right, representative image of Npy^Arc^ hM4Di-mCherry expression and Trh^Arc^ ChR2-eYFP expression and caudal fiber implant location. **B,** Average post-fast food intake during Trh^Arc^-ChR2 concurrent photostimulation and/or Npy^Arc^ inhibition (n=8 for Trh^Arc^;Npy mice, males and females, repeated-measures two-way ANOVA, time x condition: F (9, 84) = 9.75, p<0.0001, Tukey’s multiple comparisons). **C,** Overnight food intake following acute liraglutide injection at baseline (pre-TeNT) and post-TeNT in Trh^Arc^-TeNT and wildtype (WT) littermates bilaterally injected with Cre-inducible AAV-EGFP-2a-TeNT (n=11 for Trh^Arc^-TeNT, n=11 for WT-TeNT,, males and females, RM two-way ANOVA, Time x Condition: F (1, 20) = 12.97, p<0.002, Tukey’s multiple comparisons). **D,** Body weight change over 1 week of daily liraglutide administration in Trh^Arc^-TeNT and wildtype (WT) littermates bilaterally injected with Cre-inducible AAV-EGFP-2a-TeNT (n=9 for Trh^Arc^-TeNT, n=8 for WT-TeNT, males and females). **E,** Daily kcal consumption over 1 week of daily liraglutide administration in Trh^Arc^-TeNT and wildtype (WT) littermates bilaterally injected with Cre-inducible AAV-EGFP-2a-TeNT (n=9 for Trh^Arc^-TeNT, n=8 for WT-TeNT, males and females). **F,** Average body weight change over time in Trh^Arc^-TeNT and wildtype (WT) littermates bilaterally injected with Cre-inducible AAV-EGFP-2a-TeNT (n=17 for Trh^Arc^-TeNT, n=17 for WT-TeNT, males and females). **G,** Average weekly chow intake over time in Trh^Arc^-TeNT and wildtype (WT) littermates bilaterally injected with Cre-inducible AAV-EGFP-2a-TeNT (n=17 for Trh^Arc^-TeNT, n=17 for WT-TeNT, males and females).

## Notes

### Competing Interest Statement

The authors have declared no competing interest.

### Summary of Updates

We revised our manuscript to include many new experimental findings and discussion points, including: 1. Trh+ Arc neurons mediate roughly half of the body weight loss from chronic treatment with the GLP-1R agonist liraglutide, revealing a more substantial role for these neurons then we previously observed with acute liraglutide treatment; 2. Trh+ Arc neurons inhibit feeding through AgRP neurons, since inhibiting AgRP neurons while activating Trh+ Arc neurons does not decrease feeding more than either manipulation alone; 3. Trh+ Arc neurons express Glp1r transcripts, are distinct from AgRP neurons, and are activated by feeding; 4. The Arc and dorsomedial hypothalamus are enriched with Glp1r+ afferents to AgRP neurons; 5. The pattern we observe of neurons labeled with monosynaptic rabies via AgRP neurons is consistent with previous studies; 6. AgRP neurons are still activated (Fos+) by fasting even after infection with rabies virus; and 7. Transferring cell-type and brain region-labels from a more comprehensive reference atlas, the mouse HypoMap, improves precision at identifying molecular and anatomical cell types among rabies-infected cells.

